# Multifaceted fusion defects converge on altered mitochondrial distribution and increased mutant mitofusin level in CMT2A models

**DOI:** 10.1101/2025.03.14.642792

**Authors:** Olena O. Varuk, Amandine Ruiz, Celia Orengo, Amelie Cozzolino, Paul de Boissier, Aïcha Aouane, Thomas Rival

## Abstract

Charcot-Marie-Tooth type 2A neuropathy (CMT2A) is caused by a hundred different missense mutations in the mitofusin MFN2, a mitochondrial fusion factor. It is well established that different CMT2A alleles expressed in mitofusin deficient cells can exhibit varying levels of fusion activity. However, most CMT2A mutations are dominant. Therefore, to picture the complexity of CMT2A pathogenesis, it is crucial to define whether and how different fusion activity levels translate into different dominant properties. To explore the range of dominant properties among CMT2A alleles, we modelled in drosophila fourteen missense mutations reported in patients. We studied alleles from different mutation hotspots identified in MFN2 protein structure, alleles causing severe or mild forms of CMT2A, or alleles with different level of dominance in patients. Mitochondrial architecture analysis revealed a wide spectrum of composite dominant effects of variable expressivity. Fusion-inhibition and aggregation properties were found in mutants affecting catalytic motifs and hydrophobic cores, and exacerbation of mitochondrial fusion in mutants targeting domain interface. In these mutant categories, mitochondrial cytoplasmic distribution was disturbed to varying degrees that strongly correlate with neurotoxicity levels. Inhomogeneity of mitochondrial repartition was indeed associated with decreased endoplasmic reticulum-mitochondrion overlap, reduced mitochondrial content at neuromuscular junctions, higher locomotor deficits in flies and higher disease severity in patients. At the molecular scale, most CMT2A mutant proteins showed reduced ubiquitination and increased protein level, likely amplifying their dominant properties. Thus, CMT2A mutations induce a broad spectrum of fusion alterations converging towards altered mitochondrial distribution and mutant Mfn accumulation.

## Introduction

Mitochondria are connected and disconnected by fusion-fission cycles, a process supporting mitochondrial homeostasis (Giacomello et al., 2020). Outer and inner mitochondrial membranes (OMM, IMM) fusion is driven by mitofusin (Mfn) and OPA1 respectively (Tilokani et al., 2018). The resulting content mixing optimises mitochondrial metabolism (Chen et al., 2005), buffers mitochondrial damages (Ono et al., 2001) and supports mitochondrial replication (Silva Ramos et al., 2019). Mitochondrial fission is driven by DRP1, facilitating mitophagy (Twig et al., 2008) and mitochondrial transport (Verstreken et al., 2005). Neurons being highly dependent on mitochondria, mitochondrial fusion-fission imbalance causes neurological disorders (Bertholet et al., 2016). Mutations affecting human mitofusin 2 (MFN2) cause type 2A Charcot-Marie-Tooth disease (CMT2A), a hereditary motor and sensory axonal neuropathy (Stuppia et al., 2015). CMT2A is mostly due to dominant alleles leading to amino acid (aa) substitutions, although rare recessive forms were described (Stuppia et al., 2015, Pipis et al., 2020). Depending on the alleles, CMT2A can be early or late onset with severe to mild symptoms (Stuppia et al., 2015, Pipis et al., 2020). Several evidence suggest that CMT2A neuronal selectivity is due to low MFN1 expression in neurons (Kawalec et al., 2014). Indeed MFN1, that is partially redundant with MFN2, is well known to rescue CMT2A mutations in cell culture and *in vivo* (Detmer and Chan, 2007, Zhou et al., 2019, Shahin et al., 2023).

Mfns are large multi-domain proteins related to bacterial dynamin-like protein (BDLP) (Low and Lowe, 2006, Low et al., 2009). A GTPase domain (GD) and two helix bundles (HB1, HB2) are linked by hinges, and a short hydrophobic segment (TM) makes a OMM anchor. HB1 and HB2 display two heptad repeats, HR1 and HR2, forming a coiled coil. Resolved structures of truncated MFN1 and MFN2 (Koshiba et al., 2004, Cao et al., 2017, Yan et al., 2018, Li et al., 2019), and *in silico* modelling of MFN2 (Beresewicz et al., 2018, Rocha et al., 2018) and yeast Fzo1 (De Vecchis et al., 2019) provided insights into putative Mfns’ activity. Interactions through GD and HR2 might drive trans-oligomerisation and subsequent mitochondrial tethering (Koshiba et al., 2004, Cao et al., 2017, Li et al., 2019). At hinges, GTP hydrolysis would then drive an upward GD tilt (Yan et al., 2018, Li et al., 2019) and a scissor-like movement bringing HB1 closer to HB2 (Beresewicz et al., 2018, Rocha et al., 2018, Brandner et al., 2019). This open to a closed conformation switch would provide OMM fusion driving force (Formosa and Ryan, 2016, De Vecchis et al., 2019). In this process, HR1-HR2 interaction might negatively regulate fusion (Franco et al., 2016, Rocha et al., 2018), HR1 playing a role in lipid bilayer destabilisation (Daste et al., 2018). Mfn activity is regulated by phosphorylation, oxidation and ubiquitination (Giacomello et al., 2020). In yeast, Fzo1 oligomer disassembly is mediated by ubiquitination and subsequent proteasome-mediated degradation (Cohen et al., 2008, Alsayyah et al., 2020). In animals, several E3 ubiquitin ligases, including Parkin, ubiquitinate Mfns, in particular in mitophagy (Escobar-Henriques and Joaquim, 2019). Finally, in addition to mitochondrial fusion activity, Mfns establish contacts between mitochondria and various organelles (Gordaliza-Alaguero et al., 2019, Huo et al., 2022, Alsayyah et al., 2024).

Drosophila shares its mitochondrial fusion-fission machinery with mammals. Marf is the only ubiquitous Mfn (Hwa et al., 2002) whose neuronal depletion triggers mitochondrial fragmentation (El Fissi et al., 2018). Like mammalian Mfns, Marf is regulated by ubiquitination (Ziviani et al., 2010, Yun et al., 2014) and oxidation (Smith et al., 2019). Marf restores mitochondrial fusion in *Mfn1-Mfn2* knockout mouse cells, demonstrating the functional conservation between fly and mammalian Mfns (El Fissi et al., 2018). Marf appears closer to human MFN2 than MFN1 as the G2 motif isoleucine 108, causing MFN1’s tenfold higher GTPase activity, is replaced by a threonine in MFN2 (T129) and Marf (T170) (Li et al., 2019). In addition, Marf depletion influences ER-mitochondrion interactions (Debattisti et al., 2014, Sandoval et al., 2014), a MFN2-specific function.

CMT2A mutations target residues in different MFN2 domains (Cartoni and Martinou, 2009) and might affect mitochondrial fusion in different ways. Consistently, several comparative studies revealed that different CMT2A alleles expressed in Mfn knockout cells can support varying levels of mitochondrial fusion (Detmer and Chan, 2007, Barsa et al., 2025, Joaquim et al., 2025). However, in the absence of endogenous MFN2, this did not tell whether and how these differences in terms of intrinsic activity translate into different dominant properties. To be in capacity to compare, among a large allelic series and in an *in vivo* context, potentially complex dominant properties, we chose drosophila as a model system. Flies present several advantages. The expression of a single mitofusin that approaches the situation in human neurons where MFN2 is preponderant. The accessibility of motor neurons that allows high resolution microscopy of mitochondria. Controlled transcription level and genetic background, and targeted expression in motor neurons. The possibility to correlate mitochondrial alterations with consequences at the organism scale, such as locomotor defects. Hence, we modelled and studied in flies fourteen CMT2A alleles, which represents the largest number of mutations ever compared so far, revealing an unanticipated broad spectrum of dominant effects.

## Results

### CMT2A mutations concentrated into structural hotspots

To highlight mutations hotspots in MFN2, we generated protein structure models based on BDLP closed and open conformations (Low and Lowe, 2006, Low et al., 2009), onto which we mapped CMT2A mutations (Figure 1A,B). From the literature, we listed ninety-four MFN2 residues mutated in CMT2A: 56% in GD, 22% in HB1, 11% in HB2, 7% in the hinges and 2% in TM (Figure 1A,B). Mutation frequency heat maps showed mutation enrichment in specific structural motifs (Figure 1C-E; Figure S1A). Secondary structures at the GD front were more affected than those at the back, particularly G1, G2, G3 and G4 catalytic motifs, and α1G, α2G, α3G and α4G helices (Figure 1C-E). Mutations affecting these regions may affect GTPase activity and oligomerisation as the homologous secondary structures form the dimer interfaces of MFN1 and BDLP (Figure S1B-E). The α3G C-terminal part and adjacent loop could form a GD-HB1 interface in MFN2 open conformation as observed in MFN1 and BDLP (Figure 1C; Figure S1B,D), whereas α2G and α3G would be at GD-HB2 interface in MFN2 closed conformation as for BDLP (Figure 1D; Figure S1E). Hence, mutations in these regions could affect the transition between open and closed conformations. In HB1 and HB2, CMT2A mutations concentrated in α3H1, α4H1 and α7H2 (Figure 1C,D) that are part of HR1 and HR2 involved in oligomerisation and fusion activity. Finally, the hinges that support Mfn conformational changes are largely affected (Figure 1C,D). Between GD and HB1, residues R94, R95, T356, K357 and S352 are mutated (Figure 1A). Homologous MFN1 residues are all involved in hinge 2 bending supporting GD movements upon GTP binding and hydrolysis (Figure S1B,C). CMT2A mutations also affected hinge 1 (Figure 1A,C,D), a region between HB1 and HB2 involved in open to closed conformational transition in BDLP (Figure S1D,E). Hence, CMT2A mutations target regions of functional importance and may lead to different molecular outcomes.

**Figure 1.**
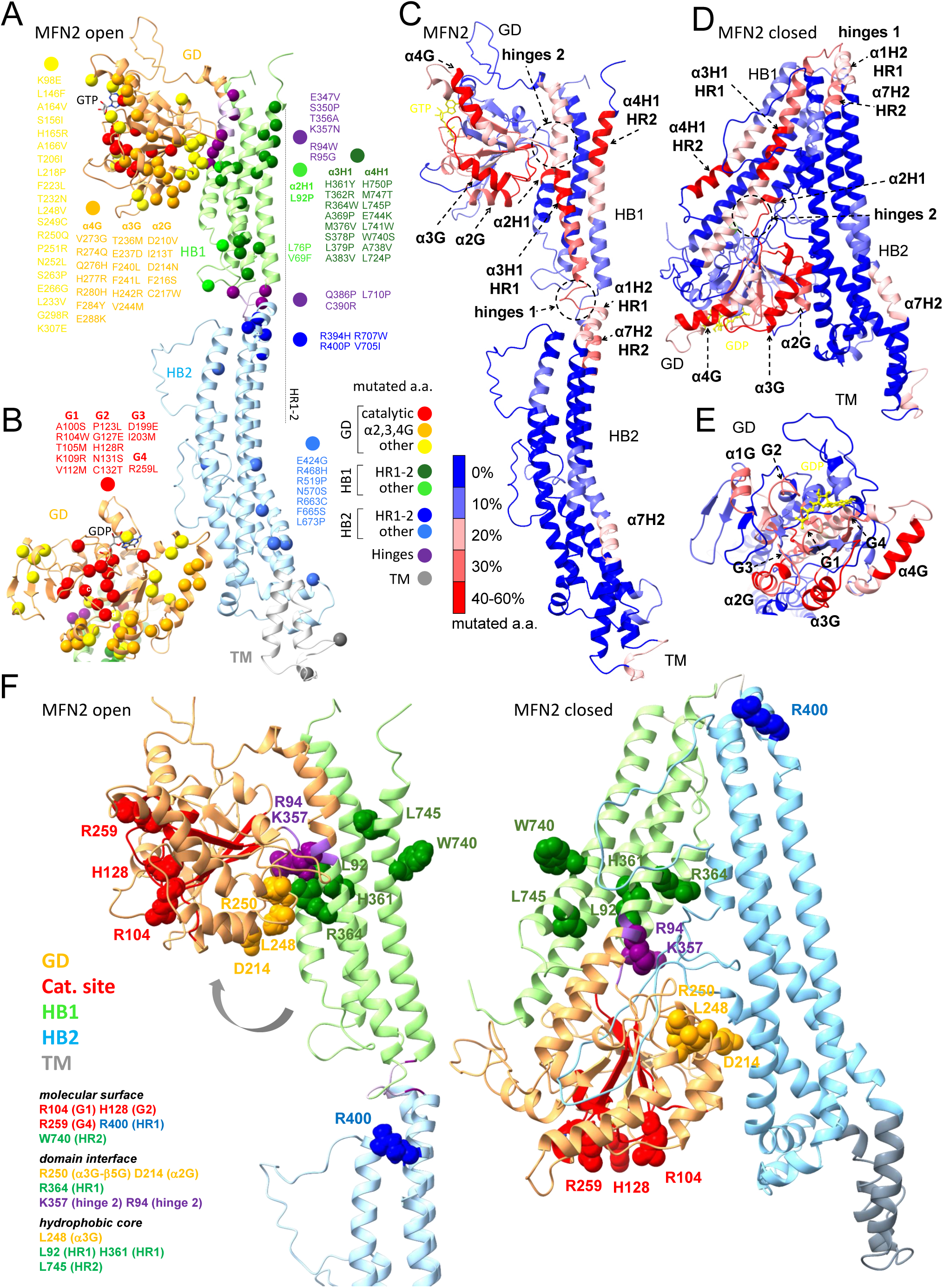
Hotspots of CMT2A mutations in MFN2. **(A)** Model of human MFN2 structure predicted by homology modelling based on BDLP (aa 1-23 removed). Spheres: carbon α of the residues mutated in CMT2A. **(B)** Highlight on the GD (front view) from MFN2 protein structure model. **(C)** Model of open MFN2 structure (based on BDLP). Colour coding: proportion of aa mutated in CMT2A within a 10 aa interval. Arrows indicate the most affected secondary structures. **(D)** CMT2A mutation frequency in MFN2 closed conformation (model based on BDLP). **(E)** CMT2A mutation frequency in MFN2 GD (front view). **(F)** MFN2 open (left) and closed (right) conformation models based on BDLP showing the residues mutated in CMT2A that were selected for our study in flies. The spheres represent the different atoms of the lateral chains (excluding hydrogen atoms). Colour code refers to MFN2 domains. Putative localisation of the selected CMT2A residues is based on their lateral chain orientation and environment.

### Modelling CMT2A allele diversity in flies

To compare CMT2A alleles from different hotspots, we mimicked them in drosophila. Marf and MFN2 share 45.5% of identical aa and 15.2% with similar properties (Figure S2A,B). When only considering CMT2A-related residues, identity and similarity reached 60.6% and 13.8% respectively (Figure S2A,B), reinforcing the idea that CMT2A-related residues are part of motifs critical for structure-function relationships. For our study in flies, we selected fourteen CMT2A aa substitutions, affecting conserved residues from different mutation hotspots: R104W, H128R and R259L for G1, G2 and G4 ; D214N for α2G ; L248V and R250Q for α3G-β5G ; R94W and K357N for hinge 2 ; L92P for α2H ; H361Y, R364W and R400P for HR1 ; W740S and L745P for HR2 (Figure 1F). We included aa at the molecular surface, at the interface between two domains, or buried within a domain (Figure 1F). In selecting these residues, we checked that their localisation relative to protein domains and their lateral chain orientation were conserved between Marf models and resolved structures of truncated MFN2 and MFN1 (Yan et al., 2018, Li et al., 2019). From there, a ‘d’ prefix refers to Marf residues (Marf, MFN2, MFN1 correspondence: Figure S2C). As for human MFN2, dR104, dH128 and dR259 localised in G1, G2 and G4 (Figure S3D; Figure S4A) and were at dimer interface, with only dR259 forming hydrogen bonds as for its MFN1 counterpart (Figure S4B). Surface orientation of dW740, that is part of HR2, was also conserved (Figure S3E; Figure S4C,E). In contrast to their orientation in the GDP-bound state (Figure S4C), dR94, dR250, dK357 and dR364 lateral chains were at the GD-HB1 interface in the GTP-bound state as for human Mfns (Figure S3C; Figure S4D). Of note, Marf dD214 and dR364 and MFN2 D214 and R364 similarly faced HB2 in closed conformation models (Figure 1F; Figure S3B). GD residue dL248 and HB1 residues dL92, dH361 and dL745 associated with hydrophobic cores as for their MFN2 equivalents (Figure S3E; Figure S4E,F). Finally, dR400 and R400 residues were predicted at HB2 surface in both Marf and MFN2 models (Figure 1F; Figure S3F). In addition to structural considerations, we explored differences in pathogenicity, by studying nine severe early onset alleles (L92P, R94W, R104W, H128R, K357N, H361Y, R364W, R400P) and five mild late onset alleles (D214N, R250Q, R259L, W740S, L745P) (Stuppia et al., 2015). Finally, we include four alleles which implication is CMT2A was classified as “uncertain” (https://www.ncbi.nlm.nih.gov/clinvar/) (D214N, R250Q, R400P, L745P), two of which being proposed to be either recessive or semi-dominant mutations (D214N, R250Q) (Nicholson et al., 2008, Piscosquito et al., 2015, Tomaselli et al., 2016). We introduced these fourteen aa substitutions into the *Marf* coding sequence, inserted all CMT2A transgenes (*Marf ^CMT2A^*) at the same *locus* and expressed them in neurons using UAS/GAL4 system (Figure S5A). All *Marf* constructs were tagged with HA. A control UAS *Marf* wild-type transgene (*Marf ^CTRL^*) was similarly generated providing an expression equivalent to 1.5 times that of endogenous Marf (Figure S5B,C).

### CMT2A alleles form four phenotypic groups with composite features

Using high-resolution confocal microscopy, we imaged for each CMT2A mutant the entire mitochondrial network of motor neurons in living larval central nervous system (CNS). To reveal mutant dominant properties, we analysed mitochondria in the presence of endogenous Marf. The resulting phenotypes were complex and composite. Therefore, we scored three categories of mitochondrial shape (fragmented, filamentous and enlarged) and two modes of interaction (net-like and clustered). In parallel, we evaluated the fusion capacity of each mutant in a *Marf* knockout background (*Marf ^KO^*) by measuring mitochondrial tubule length. These data were analysed by hierarchical clustering allowing us to classify our mutants into four phenotypic groups and to evaluate their phenotypic distance from wild-type, *Marf ^KO^* and *Marf ^CTRL^* flies (Figure 2A). These mutant groups were named Grape, Tangle, Bulb and Peanut according to their mitochondrial shape (Figure 2B).

**Figure 2.**
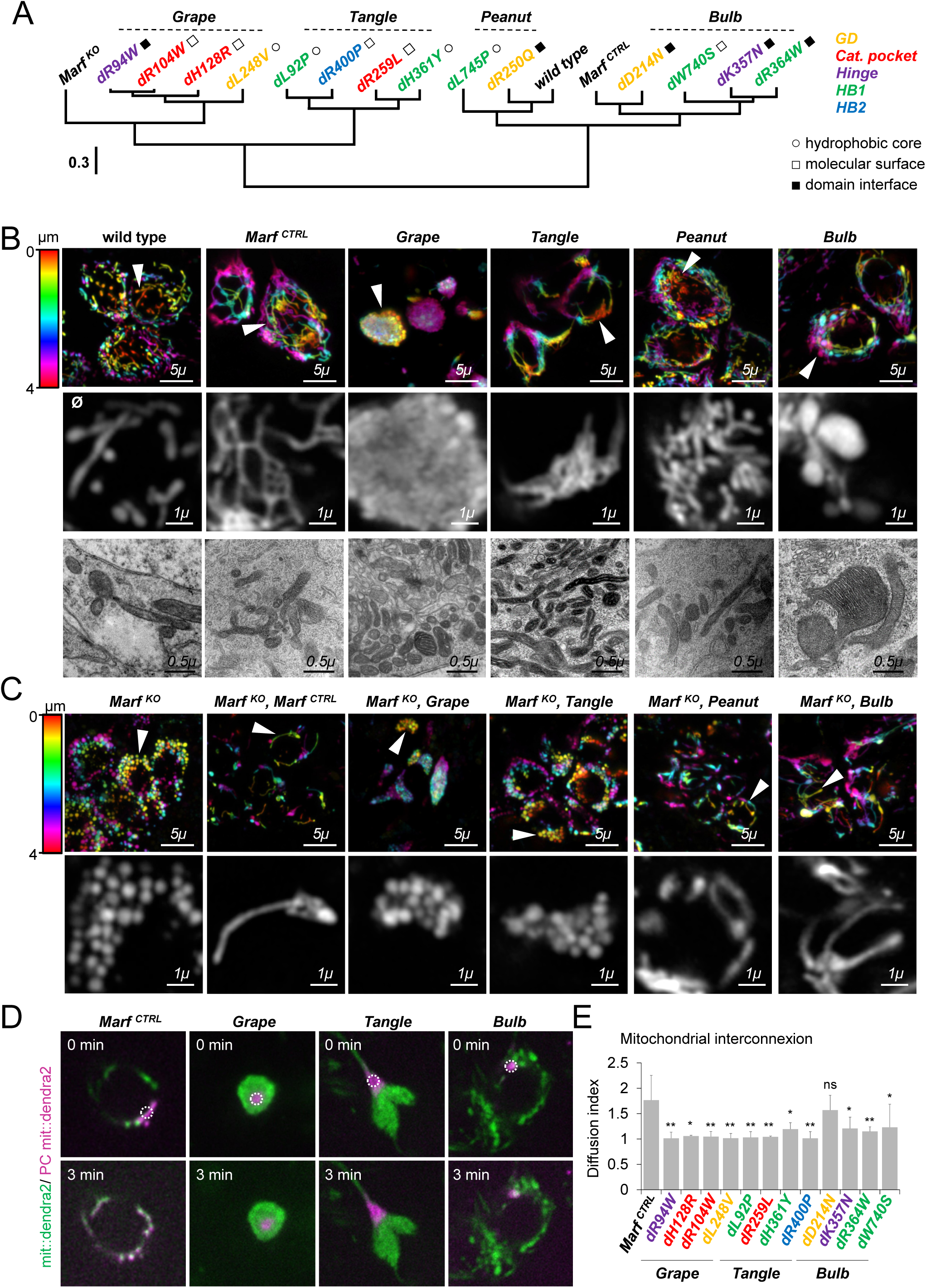
CMT2A alleles differently alter mitochondrial shape and organisation. **(A)** Dendrogram based on hierarchical clustering of mitochondrial morphology in wild-type and *Marf ^KO^* backgrounds. Hydrophobic core: aa lateral chain pointing toward the inside and associated with a network of hydrophobic residues. Molecular surface: aa lateral chain pointing toward the surface of the molecule. Domain interface: aa lateral chain in a given domain pointing toward another domain. **(B)** Confocal Airyscan image Z-projections (20 slices) of mitochondrial networks from living motor neurons (ventral nerve cord, dorsal clusters) of third instar larva (*OK37l-GAL4, UAS-mit::dendra2)*. Colour code: Z-depth in dorso-ventral axis. Wild type: flies with no *UAS-Marf* transgene. *Marf ^CTRL^*: *UAS-HA::Marf ^WT^*. Grape (*dRl04W*), Tangle (*dL92P*), Bulb (*dK357N*), Peanut (*dL745P*): *UAS-HA::Marf ^CMT2A^*. Arrows highlight mitochondrial morphological traits shown at higher magnification in the optic slices of bottom panel: tubular and fragmented (wild type and Peanut), net-like (*Marf ^CTRL^*), clustered and fragmented (Grape), clustered and filamentous (Tangle), enlarged (Bulb). TEM mitochondria high magnification views in motor neurons (dorsal clusters) from ventral nerve cord of third instar larvae (*OK37l-GAL4, UAS-mit::dendra2*). **(C)** Confocal Airyscan Z-projections (20 slices) of mitochondria from motor neurons (ventral nerve cord, dorsal clusters) of male *Marf ^KO^*/*Y, 3xP3-mCherry* ; *OK37l-GAL4, UAS-mit::dendra2*. *Marf ^CTRL^*: *UAS-HA::Marf ^WT^*. Grape (*dRl04W*), Tangle (*dL92P*), Bulb (*dK357N*), Peanut (*dR250Q*): *UAS-HA::Marf ^CMT2A^*. Colour code: Z-depth, dorso-ventral axis. Arrows: regions shown at high magnification in the bottom panels. **(D)** Mit::dendra2 photoconversion (non-photoconverted in green and photoconverted in magenta) in living motor neurons (ventral nerve cord, dorsal clusters) from *OK37l-GAL4, 2x UAS-mit::dendra2IUAS-Marf* third instar larvae. Pictures show first and last snap shots of a 3 min movie. White circle: position of the photoconvertion beam. **(E)** Mean diffusion index corresponding to the ratio of final area containing photoconverted dendra2 over initial photoconverted area +/-SD. Mann-Whitney (MW) U test: p-value <0.05/*, p-value <0.01/**, non-significant/ns.

### CMT2A mutants alter Mfn activity in different manners

Consistent with a balanced fusion-fission equilibrium, wild-type motor neurons contained both filamentous and fragmented mitochondria (Figure 2B, 3A; Figure S6A). Expression of *Marf ^CTRL^* transgene decreased mitochondrial fragmentation and triggered mitochondrial filament fusion into net-like structures (Figure 2B, 3A; Figure S6A). On the opposite, *Marf ^KO^* triggered a mitochondrial fragmentation rescued by *Marf ^CTRL^* expression (Figure 2C; Figure 3B; Figure S7A). In Grape mutants, including *dR94W*, *dRl04W* and *dHl28R*, motor neurons contained one or two large and spherical clusters of fragmented mitochondria that often showed IMM swirls or cristae loss (Figure 2B; Figure 3A; Figure S6B). Photoconversion experiments using mit::dendra2 showed that, despite being packed, mitochondria were not connected (Figure 2D,E). Grape alleles did not restore tubules in *Marf ^KO^* animals, showing fragmented and clustered mitochondria (Figure 2C; Figure 3B; Figure S7B). Taken together, these data demonstrate that Grape mutations are strong dominant negatives (DN) that do not affect Mfn tethering but abolish wild-type Mfn fusion. Consistently, Grape mutants were the closest to *Marf ^KO^* (Figure 2A). Within Tangle alleles *dL92P*, *dL248V*, *dR259L* and *dH361Y*, we observed elongated clusters of packed mitochondrial tubules (Figure 2B; Figure 3A; Figure S6C) that did not form an interconnected network (Figure 2D,E). In *Marf^KO^* neurons, Tangle mitochondria remained largely fragmented although few very short tubules were seen (Figure 2C; Figure 3B; Figure S7C). Hierarchical clustering placed Tangle close to Grape and *Marf ^KO^* (Figure 2A). In conclusion, Tangle mutants act as weak DN that cluster mitochondria without abolishing endogenous Marf fusion activity. In Peanut mutants *dR250Q* and *dL745P*, mitochondria essentially conserved their wild-type shape (Figure 2B; Figure 3A; Figure S6D). Because some *dR250Q* neurons contained net-like mitochondria (Figure 3A), we quantified mitochondrial length and branching to possibly detect subtle activity. This revealed that *dR250Q* and, to a lesser extent, *dL745P* have a residual fusion capacity (Figure S8A-D). Consistently, *dR250Q* formed mitochondrial tubules when expressed in *Marf ^KO^*, whereas *dL745P* only showed scarce short elongated mitochondria (Figure 2C; Figure 3B; Figure S7D). According to their phenotype, Peanut alleles segregated with wild-type, *dR250Q* being closer than *dL745P* (Figure 2A). Finally, Bulb mutants, including *dD214N*, *dK357N*, *dR364W* and *dW740S*, were placed by hierarchical clustering close to *Marf ^CTRL^* (Figure 2A). These alleles contained enlarged mitochondria with normal cristae and no swelling (Figure 2B; Figure 3A; Figure S6E) and restored mitochondrial tubules in *Marf ^KO^* (Figure 2C; Figure 3B; Figure S7E), showing that Bulb alleles somehow favour fusion. Paradoxically, the hyperfused bulb-like mitochondria were generally disconnected to the mitochondrial network, except for *dD214N* (Figure 2D,E). Regarding structure-function relationship, there was no correlation between the mitochondrial phenotype associated with a particular mutation and the specific protein domain affected by that mutation (Figure 2A). However, most fusion-deficient alleles affected catalytic motifs and hydrophobic cores whereas most of those that promote fusion affected interfaces between domains (Figure 2A). Of note, the alleles that clustered closer to fusion-competent controls (*dD214N*, *dR250Q* and *dL745P*) cause mild CMT2A symptoms in patients (Figure 2A). In conclusion, CMT2A mutations have different consequences on Mfn activity in flies.

**Figure 3.**
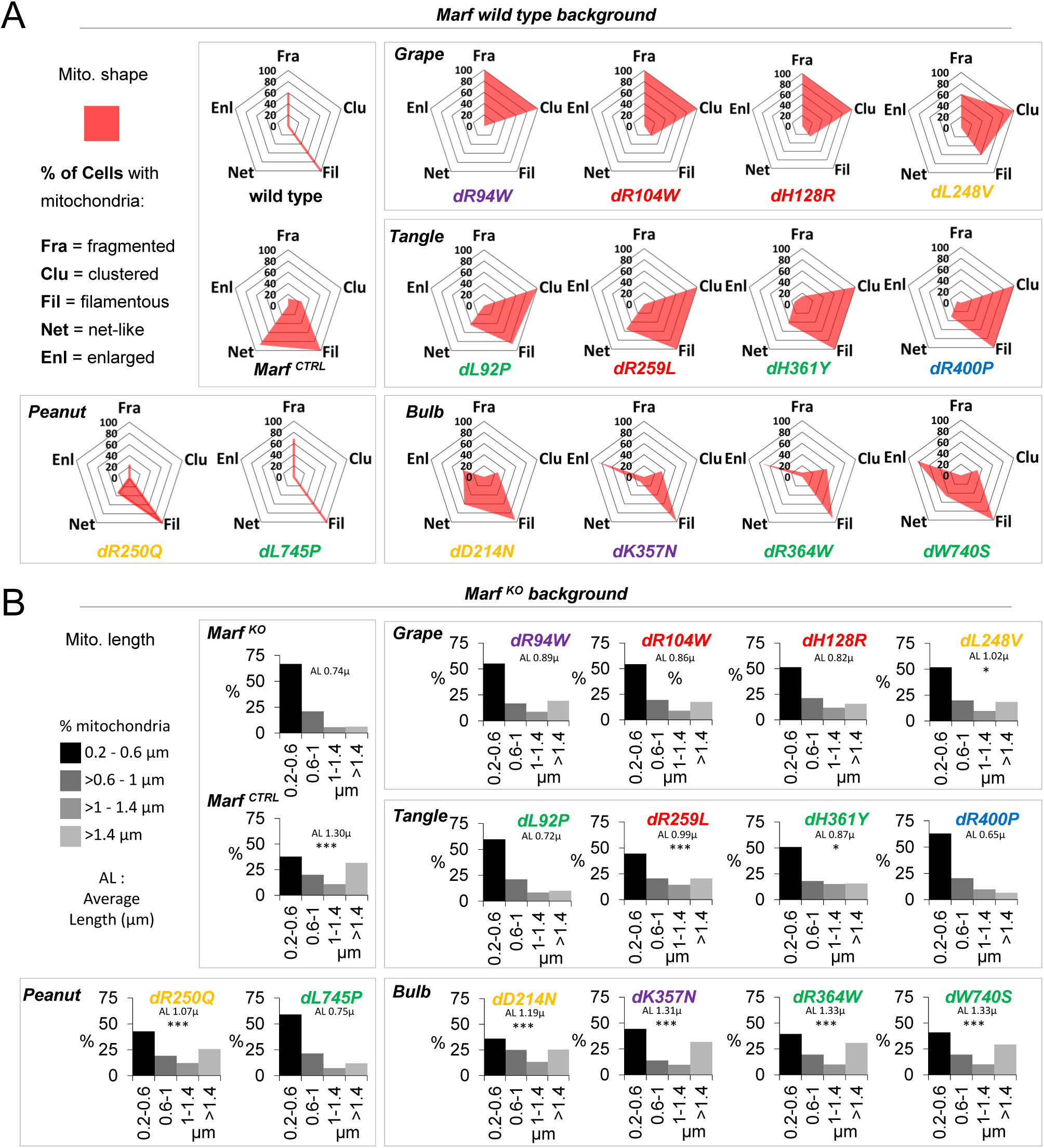
Quantitative analysis of mitochondrial networks in CMT2A mutants. **(A)** Quantifications of mitochondrial morphology in larval motor neurons based on confocal Airyscan image analysis. A given cell can contain one or several morphological traits: fragmented, clustered, filamentous, net-like, enlarged. At least three individuals analysed per conditions. On average twelve motor neurons scored per conditions (minimum: seven cells). **(B)** Distribution of mitochondria (% of total mitochondria) in *Marf ^KO^* larvae according to their length (µm). AL: average length (µm). At least 3 individuals analysed with an average total of 338 mitochondria (minimum: 167). MW U test versus *Marf ^KO^* p-value <0.05/*, p-value <0.001/***. Colour code correspond to protein domains (see Figure 1F).

### Neurotoxicity level in CMT2A flies correlates with disease severity in CMT2A patients

To determine whether the mutant mitochondrial phenotypes translate into neurotoxicity, we analysed developmental lethality, photoreceptor neurodegeneration and locomotor performance in flies upon neuron-specific expression of CMT2A mutations. Of note, startle-induced locomotion was not dependent on vision (Figure S9). Consistent with their strong DN activity, Grape mutants were the most severely affected. Only few escapers reached adulthood exhibiting high photoreceptor loss and very poor locomotor ability (Figure 4A-C). Tangle and Bulb phenotype expressivity varied between alleles with higher development lethality, stronger neurodegeneration and lower locomotion in *dL248V, dL92P* and *dH361Y* for Tangle, and *dR364W* and *dK357N* for Bulb (Figure 4A-C). Finally, Peanut mutants had no obvious toxicity (except a weak degeneration in *dR250Q*) consistent with their mitochondrial phenotype (Figure 4A-C). Importantly for disease relevance, mutants with weak or no detectable neurotoxicity in flies correspond to human alleles causing moderate CMT2A (*dR259L, dW740S* and *dL745P*) or acting as semi-dominant (*dD214N*, *dR250Q*). In conclusion, whereas Grape and Peanut phenotypes correlate with toxicity level, Tangle and Bulb alterations have variable outcomes.

**Figure 4.**
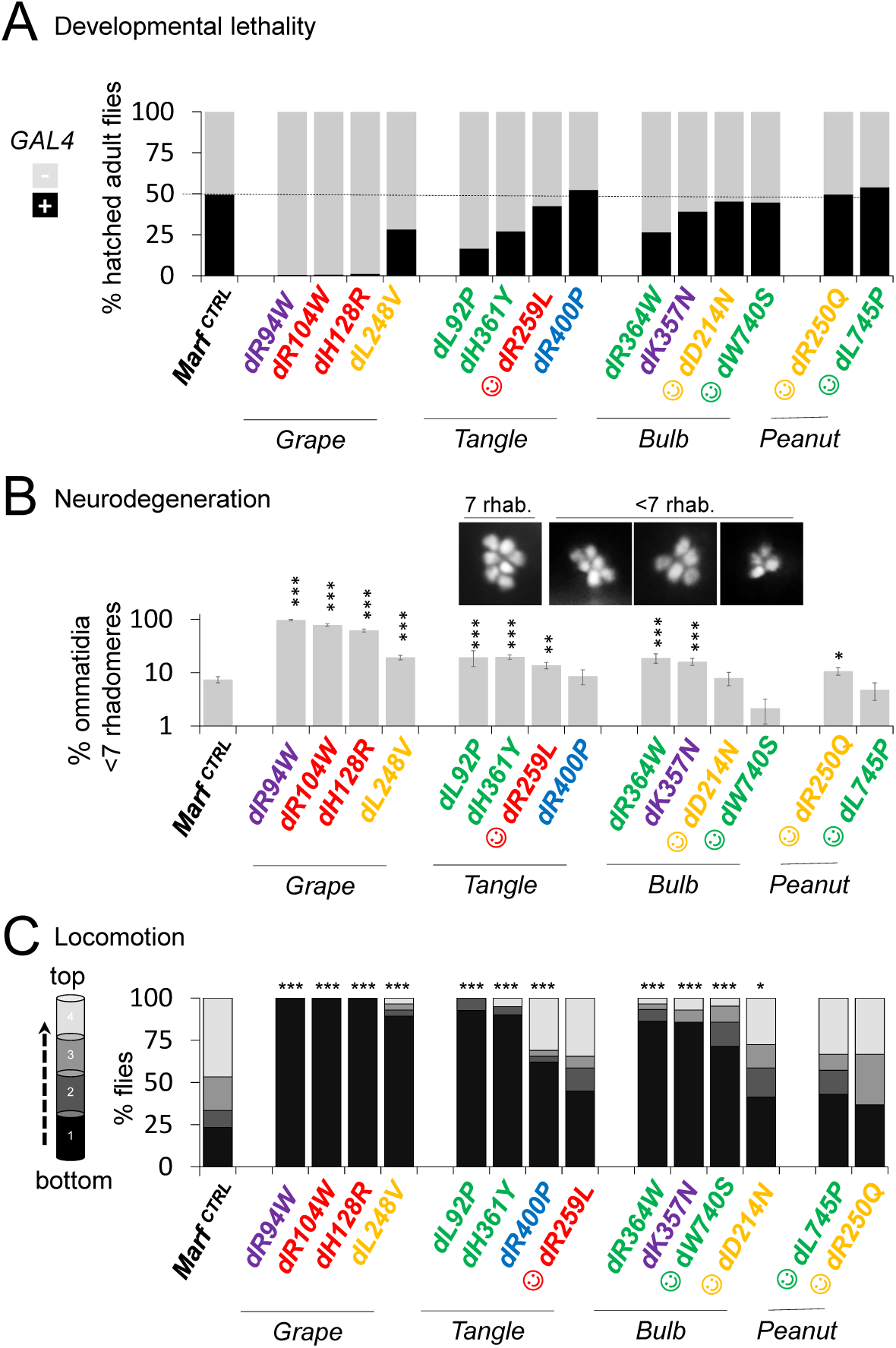
Varying degrees of neurotoxicity between CMT2A alleles. **(A)** Frequency of adult fly emergence *UAS-Marf* (*Marf ^CTRL^* or *Marf ^CMT2A^*) flies with or without *elav-GAL4* driver (average of 380 flies analysed per condition, 3 independent experiments) ranked in each allelic group according to decreasing phenotypic strength. Dashed line: expected Mendelian frequency. ☺: moderate CMT2A or semi-dominant. **(B)** Average proportion of ommatidia containing less than seven visible photoreceptor rhabdomeres +/-SEM in the retina of one week old *elav-GAL4 ; UAS-mit::dendra2IUAS-Marf* (*Marf ^CTRL^* or *Marf ^CMT2A^*) adult flies ranked by allelic groups according to decreasing phenotypic strength. Images: ommatidia with seven or less than seven photoreceptors. On average 175 ommatidia analysed from eight individuals per condition. MW U test: p-value <0.05/*, p-value <0.01/**, p value <0.001/***. ☺: moderate CMT2A or semi-dominant. **(C)** Average locomotor performance (startle-induced negative geotaxis) of ten days old *elav-GAL4 ; UAS-mit::dendra2IUAS-Marf* (*Marf ^CTRL^* or *Marf ^CMT2A^*) adult flies. Histogram shows the proportion of flies according to the height they have reached (1: 0-15 mm, 2: >15-30 mm, 3: >30-45mm, 4: >45mm). Mean of 3-4 independent races involving 20-30 flies. MW U test: p-value <0.05/*, p-value <0.01/**, p-value <0.001/***, non-significant/ns. ☺: moderate CMT2A or semi-dominant. Colour code correspond to protein domains (see Figure 1F).

### Neurotoxicity correlates with altered mitochondrial distribution

Consistent with the above observations, only two mitochondrial traits significantly correlated with decreased locomotor performance: the presence of clustered mitochondria and the loss of tubular ones (Table S1). To understand the variable neurotoxicity level within our allelic series, we searched for a common factor which strength could vary between alleles. One obvious parameter was the distribution of mitochondria within motor neurons. Indeed, distribution could be altered both by clustering in Grape and Tangle and by hyperfusion in Bulb. Quantification revealed that mitochondria were homogeneously distributed in wild-type as well as in Peanut mutants, whereas they had a focal localisation in Grape mutants (Figure 5A,B; Figure S10A). Mitochondrial distribution inhomogeneity was intermediate for Tangle and Bulb alleles, and varied between mutants (Figure 5A,B; Figure S10A). We then plotted for each allele mitochondrial distribution index to locomotor performance and found a significant correlation (Figure 5I). Hence, the more mitochondrial distribution was altered, the higher the neurotoxicity. We then investigated whether altering mitochondrial distribution affects ER organisation and interaction with mitochondria (Figure 5C; Figure S10B). Unlike mitochondria, ER had a normal distribution in CMT2A neurons (Figure 5D). Hence, the asymmetric distribution between ER and mitochondria led to decreased level of ER signal overlapping mitochondria in the most toxic CMT2A alleles, including all Grape mutants, Tangle mutants *dL248V, dL92P* and *dH361Y* and Bulb mutants *dR364W* and *dK357N* (Figure 5E). Grape mitochondria were less associated to the ER that was abnormally fragmented within mitochondrial clusters (Figure 5F; Figure S10B). In contrast, the overlap of mitochondria with ER tubules was unchanged for Tangle and Bulb mutants and increased in Peanut mutants (Figure 5F; Figure S10B). Because synaptic mitochondria are transported from neuronal soma, we tested whether altered distribution in cell bodies could reduce mitochondria at neuromuscular junctions (NMJ). Most CMT2A alleles had decreased mitochondrial content at NMJ, the most severe depletion being observed for the most toxic mutants (Figure 5G,H; Figure S10C). Statistical analysis revealed that reduced ER-mitochondrion overlap and NMJ mitochondria correlated with inhomogeneous mitochondrial distribution and locomotor deficit (Table S1). In conclusion, the extent to which a CMT2A allele is toxic in flies is proportional to the extent of which it alters mitochondrial distribution.

**Figure 5.**
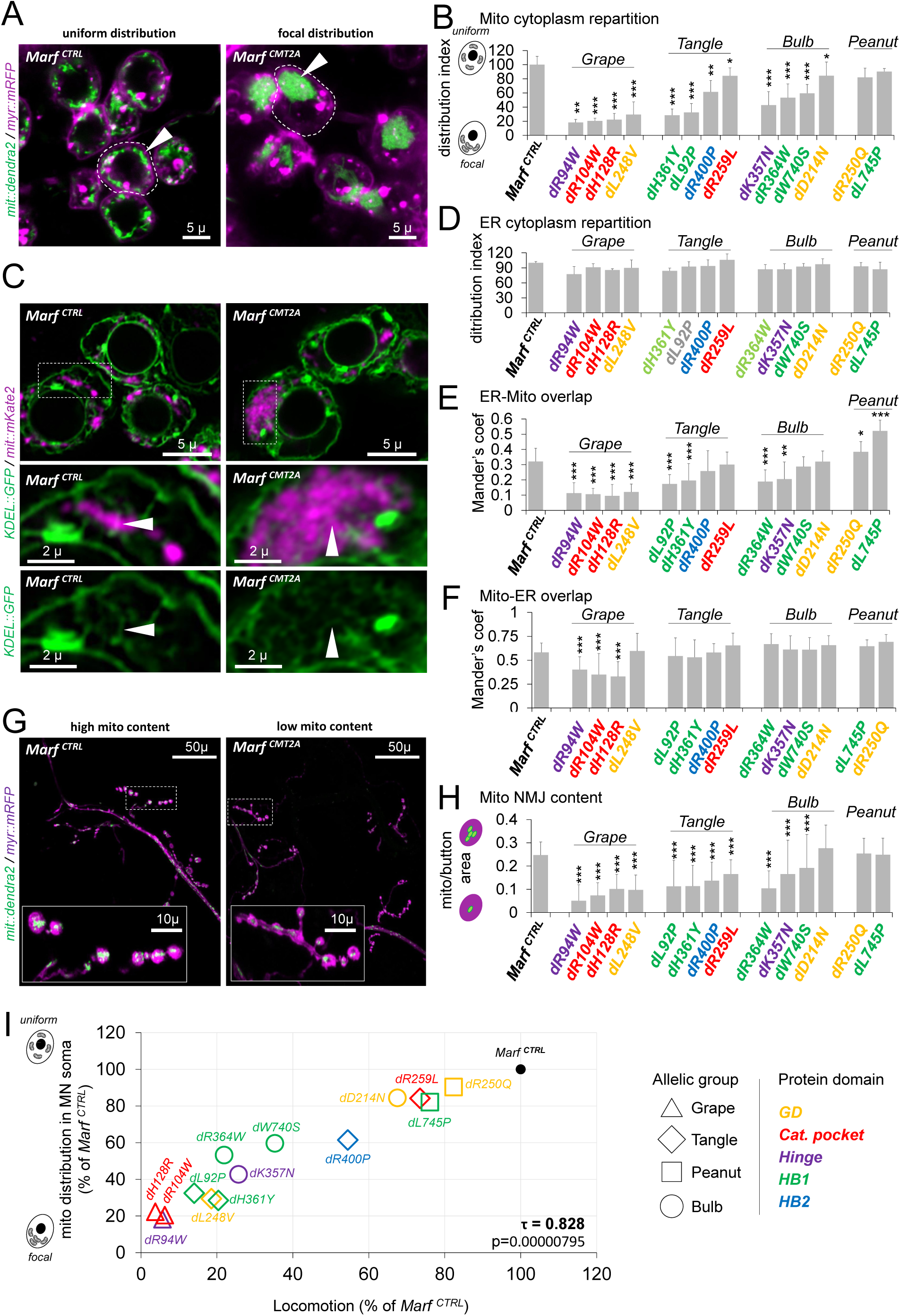
Alterations of mitochondrial distribution correlates with neurotoxicity. **(A)** Confocal microscopy of living motor neurons showing mitochondria and plasma membrane in *OK37l-GAL4, UAS-mit::dendra2 ; UAS-myr::mRFP* third instar larva expressing *Marf ^CTRL^* and *Marf ^CMT2A^ (dL92P*). **(B)** Mean mitochondrial distribution index +/-SD. Variance of all mitochondrial pixel coordinates (*x*, *y*) was analysed on average on nine cell bodies. MW U test: p-value <0.05/*, p-value <0.01/**, p-value <0.001/***. **(C)** Confocal Airyscan microscopy images of living motor neurons showing mitochondria and ER in *OK37l-GAL4, UAS-mit::mKate2, UAS-KDEL::GFP* third instar larva expressing *Marf ^CTRL^* and *Marf ^CMT2A^ (dL92P*). Magnification views: ER and mitochondria in close vicinity and overlapping (arrow heads). **(D)** Mean distribution index of endoplasmic reticulum for the different CMT2A alleles +/-SD. At least, three cell bodies analysed per condition. MW U test: not significant. **(E)** Histogram showing mean Manders’ coefficient (percent of *Marf ^CTRL^*) representing the fraction of KDEL::GFP (ER) signal overlapping with mit::mKate2 (mitochondria) staining +/-SD (sum of GFP pixel intensities with mKate2 intensity above 0 divided by the total GFP pixel intensity). Average image number analysed: 19 (minimum 14). MW U test: p-value <0.05/*, p-value <0.01/**, p-value <0.001/***. **(F)** Mean Manders’ coefficients (percent of *Marf ^CTRL^*) representing the fraction of mit::mKate2 (mitochondria) signal overlapping with KDEL::GFP (ER) staining +/-SD (sum of mKate2 pixel intensities with GFP intensity above 0 divided by the total mKate2 pixel intensity). Average image number analysed: 19 (minimum 14). MW U test: p-value <0.05/*, p-value <0.01/**, p-value <0.001/***. **(G)** Confocal Airyscan microscopy images showing mitochondria within NMJ boutons in living muscles from *OK37l-GAL4, UAS-mit::dendra2 ; UAS-myr::mRFP* third instar larva expressing *Marf ^CTRL^* and *Marf ^CMT2A^ (dL92P*). Insight: high magnification view of synaptic boutons. **(H)** Mean ratio of mitochondrial over bouton area +/-SD. Average boutons analysed: 100). MW U test: p-value <0.05/*, p-value <0.01/**, p-value <0.001/***. **(I)** Correlation plot between mitochondrial distribution in motor neuron soma (data Figure 5A) and locomotor performances (data Figure 4C) both expressed as percent of *Marf ^CTRL^*. τ: Kendall correlation coefficient. p: p-value.

### Grape and Tangle alleles are insensitive to Parkin overexpression

Searching for mechanisms that could commonly rescue Grape, Tangle and Bulb mutants, we thought that increasing Mfn turnover could help neurons to cope with defective CMT2A proteins. In flies, Marf is targeted to degradation by E3 ubiquitin ligase Parkin (Ziviani et al., 2010). Consistently, Parkin overexpression (*parkin^OE^*) in *Marf ^CTRL^* increased fragmentation and reduced fusion into net-like structures (Figure 6A,B), whereas *parkin* knockdown induced enlarged bulb-like mitochondria (Figure S11A). In Grape and Tangle mutants, *parkin^OE^* did not prevent mitochondrial clustering suggesting that the mutant proteins were resistant to Parkin-dependent degradation (Figure 6A,B; Figure S11B,C). In Tangle, *parkin^OE^* triggered a fragmentation of mitochondria that was stronger than the one observed in *Marf ^CTRL^* control (Figure 6A,B; Figure S11C). This confirmed that Tangle alleles are inactive and suggested that the mitochondrial tubules found in these mutants were generated by endogenous Marf. Similarly, *parkin^OE^* led to mitochondrial fragmentation in Peanut mutants (Figure 6A,B; Figure S11D). *Parkin^OE^* induced mitochondrial clustering in *dR250Q* (Figure 6A,B; Figure S11D), possibly reflecting a cryptic DN property revealed by wild-type Marf degradation. Consistent with these mitochondrial phenotypes, *parkin^OE^* increased developmental lethality in Grape, Tangle and Peanut mutants supporting the idea that Parkin might target wild-type but not mutant Marf (Figure 6C). In contrast, *parkin^OE^* lowered the occurrence of enlarged mitochondria in Bulb mutants (Figure 6A,B; Figure S11E), and developmental lethality was not exacerbated by Parkin except for *dK357N* (Figure 6C). Taken together, these observations demonstrated that most CMT2A fly mutants are insensitive to Parkin.

**Figure 6.**
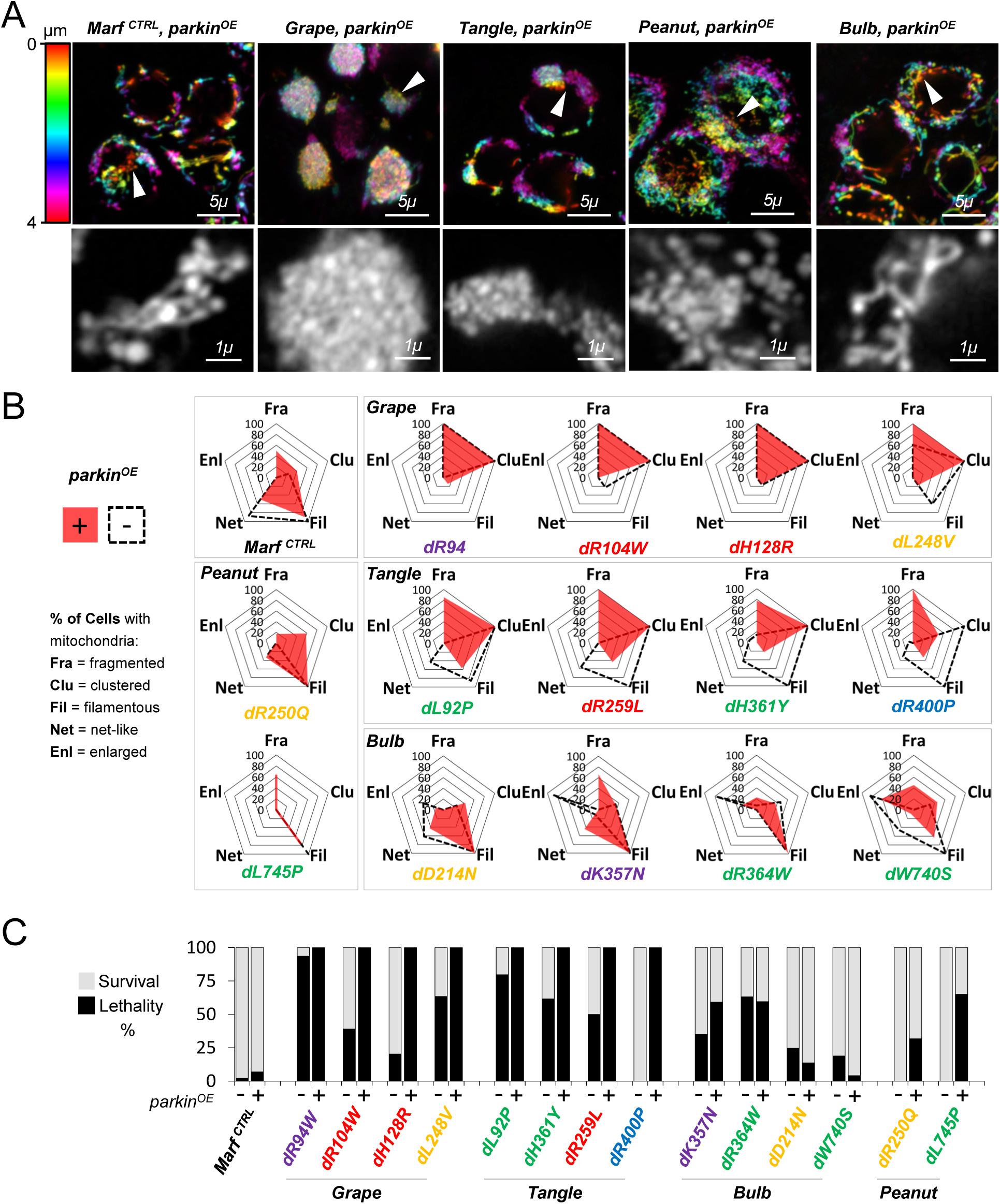
Most CMT2A alleles are insensitive to Parkin overexpression. **(A)** Confocal Airyscan images from living motor neuron of *OK37l-GAL4, UAS-mit::dendra2I UAS-parkin* (*parkin^OE^*) third instar larva expressing *Marf ^CTRL^* or *Marf ^CMT2A^*. The mitochondrial structures pinpointed by arrows are magnified on bottom panel images. Grape: *dR94W*. Tangle: *dL92P*. Peanut: *dL745P*. Bulb: *dR364W*. **(B)** Quantifications of mitochondrial morphology in larval motor neurons overexpressing Parkin (*parkin^OE^*) based on confocal Airyscan image stacks. A given cell can contain mitochondria with one or several morphological traits: fragmented, clustered, filamentous, net-like, enlarged. Black lines: phenotypes without *parkin^OE^*. Red surfaces: phenotype upon *parkin^OE^*. At least three individuals were analysed. Average total number of motor neurons analysed: 11 (minimum 6). **(C)** Developmental lethality with (+) or without (-) *parkin* overexpression (*parkin^OE^*). Males *elav-GAL4IY ; UAS-mit::dendra2* flies were crossed to *UAS-parkin* females combined with *UAS-Marf ^CTRL^ or UAS-Marf ^CMT2A^* and were kept at 29°C. In the resulting adult progeny, the number of males (no *elav-GAL4*) and females (with *elav-GAL4*) was determined (average of 150 flies per conditions). To minimize developmental lethality, *dR94W*, *dRl04W* and *dHl28R* were crossed to *OK37l-GAL4, UAS-mit::dendra2ICyO* and kept at 25°C. Flies with Curly and wild-type wings were scored. Colour code correspond to protein domains (see Figure 1F). The difference in percent between males and females, or Curly and non-Curly populations referred as percent survival. The percent lethality corresponds to 100 minus the percent survival.

### Reduced ubiquitination and increased protein levels in CMT2A mutant flies

We then investigated whether CMT2A mutations interfere with Marf ubiquitination. Our *Marf ^CTRL^* and *Marf ^CMT2A^* transgenes being HA-tagged, we could specifically analysed these recombinant proteins independently of endogenous Marf. We first determined Marf ubiquitination profile in *Marf ^CTRL^* flies. Anti-HA western blot against immuno-precipitated Marf revealed three main bands at 100 kDa, 110 kDa and 130 kDa (Figure 7A). Anti-Ubiquitin staining demonstrated that the 100 kDa band corresponded to native Marf whereas the 110 and 130 kDa bands were ubiquitinated forms (Figure 7A). Of note, a minor band with higher molecular weight and higher ubiquitination was also detected. Parkin was involved in the formation of the two main Marf ubiquitinated forms as they both decreased upon *parkin* knockdown (*parkin^KD^*) (Figure 7B). We next determined ubiquitinated *versus* native Marf ratios in CMT2A mutants. Ubiquitination was reduced for all CMT2A proteins except for *dD214N* (Figure 7C,D). Interestingly, the ubiquitination level of mutant proteins correlated with the fusion capacity measured in *Marf ^KO^* CMT2A flies (Table S1). This ubiquitination defect was followed by increased mutant Marf level for most CMT2A strains (Figure 7E) which significantly correlated with decreased locomotor performance as well as focal distribution and clustering of mitochondria (Table S1). In conclusion, CMT2A mutations affect Marf ubiquitination, likely potentiating their dominant properties through mutant protein accumulation.

**Figure 7.**
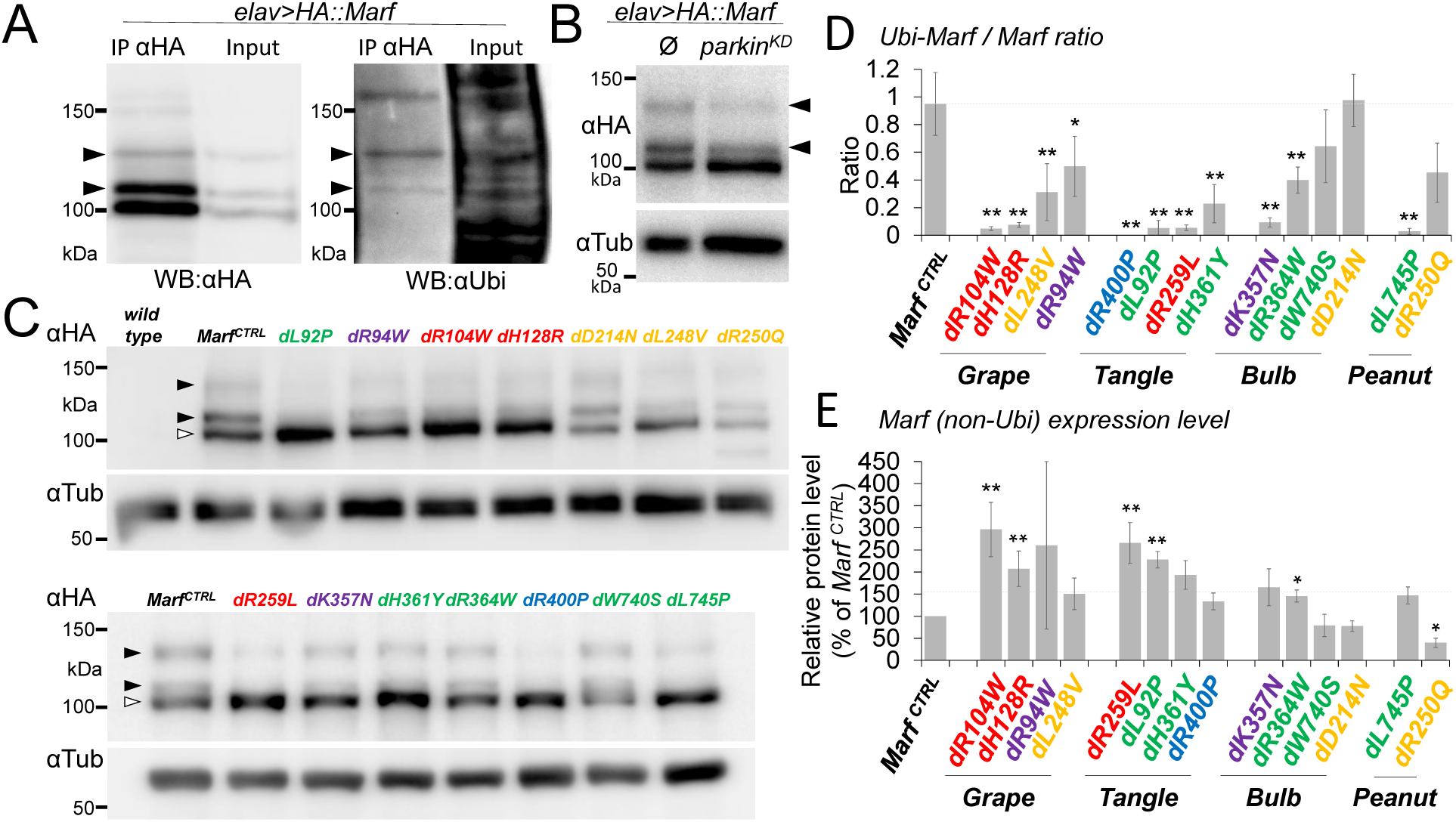
Decreased ubiquitination and increased protein level in CMT2A mutant Marf. **(A)** Anti-HA and anti-Ubiquitin western blots of immunoprecipitated wild-type HA-tagged Marf (IP) from *elav-GAL4 ; UAS-mit::dendra2IUAS-HA::Marf* (*Marf ^CTR^*^L^) third instar larva CNS. Input: protein extract before IP. Same blot first stained with anti-Ubiquitin, and following stripping, re-probed with anti-HA. Black arrows: ubiquitinated Marf. **(B)** Anti-HA western blot of proteins from *elav-GAL4 ; UAS-mit::dendra2IUAS-HA::Marf* (*Marf ^CTR^*^L^) larval CNS alone (0) or with *UAS-RNAi* against *parkin* (*parkin^KD^*). Black arrows: ubiquitinated Marf. **(C)** Anti-HA and anti-α-Tubulin western blot of proteins from CNS of *elav-GAL4 ; UAS-mit::dendra2IUAS-HA::Marf* (*Marf ^CTR^*^L^ or *Marf ^CMT2A^*) larvae. Black arrows: ubiquitinated Marf. White arrow: native Marf. **(D)** Average ratio between ubiquitinated and native Marf based on western blot signal analysis. Mean +/-SD of 3-4 experiments involving independent samples (5 CNS). MW U test: p-value <0.05/*, p-value <0.01/**. **(E)** Average native Marf level (arbitrary units) based on western signal intensity (normalisation: α-Tubulin). Mean +/-SD of 3-4 experiments involving independent samples (5 CNS). MW U test: p-value <0.05/*, p-value <0.01/**. Colour code correspond to protein domains (see Figure 1F).

## Discussion

We showed that CMT2A mutations are enriched in specific MFN2 regions, many of which previously shown to play key roles in Mfns and BDLP activity. These hotspots include GTP binding pocket motifs and GD secondary structures that form the dimer interface. In HB1, mutations accumulates in HR1 and HR2 which regulate trans-oligomerisation and membrane fusion. CMT2A mutations also strongly affect the flexible hinges supporting GD movements and HB1-HB2 closure. Finally, it is noticeable that several residues localize at the interface between GD and HB1 in the open conformation. Our observations have several general implications. For clinicians, the hotspots we defined could improve MFN2 mutation scoring: a novel variant having more chance to be pathogenic if it targets an aa from a hotspot. CMT2A hotspots could also orient structure-function studies as they highlight specific structures of interest and specific residues within these structures which analysis could improve our knowledge on Mfn molecular mechanism.

We classified our CMT2A mutants into four groups: Grape, Tangle, Peanut and Bulb. Grape mutants act as strong DN with a high aggregative property, observed even in the absence of endogenous Marf. Hence, oligomerisation capacity was retained but fusion capacity was lost. Such phenomenon could occur because catalytic pocket mutants *dHl28R* and *dRl04W* would affect GTP hydrolysis, or because hinge 2 mutant *dR94W* would compromise conformational changes triggered by GTP hydrolysis. The fact that *dL248V* had a similar, although milder, Grape phenotype could result from the alteration of GD hydrophobic core in which L248 is implicated. In *Marf ^KO^*, Tangle alleles had fragmented mitochondria that did not cluster. In contrast, they induced mitochondrial aggregation in the presence of wild-type Marf. However, unlike Grape mutants, Tangle alleles did not fully counteract wild-type Marf activity, mitochondria remaining tubular. This suggests that Tangle mutant proteins do not efficiently assemble with each other to form fusion complexes, but can interact with wild-type Marf. In that case, the resulting fusion complexes might remain locked at the docking step, causing mitochondrial clustering. Two Tangle mutants, *dL92P* and *dH361Y*, are part of a large hydrophobic network maintaining HB1 helices (Yan et al., 2018, Li et al., 2019), another one, *dR259L*, is part of G4 catalytic motif, and a third one, *dR400P*, is a HB2 surface residue. Interestingly, R259 in addition of being part of the catalytic pocket, as for R104 and H128, is involved in MFN2 dimerisation through an interaction with E230 from the opposite molecule (Yan et al., 2018, Li et al., 2019). Hence, *dR259L* by reducing the ability of mutant molecules to interfere with wild-type ones could lead to a weaker DN phenotype than *dRl04W* and *dHl28R*. Peanut mutants had reduced fusion activity, *dR250Q*, at the GD-HB1 interface, being more active than *dL745P* that is part of HB1 hydrophobic core (Li et al., 2019). In *Marf ^KO^* flies, Bulb mutants led to mitochondrial tubule formation and, in wild types, they formed enlarged bulb-shape mitochondria typically described in mammalian and fly cells deficient for DRP1 or overexpressing Mfn (Wakabayashi et al., 2009, Rival et al., 2011, Kageyama et al., 2012, El Fissi et al., 2018). Of note, these hypertrophied mitochondria were generally disconnected from the rest of the network likely interfering with global mitochondrial content mixing. In Detmer and Chan’s landmark study, four out of nine CMT2A mutations restored mitochondrial tubules in *Mfn1*-*Mfn2* double knockout cells (Detmer and Chan, 2007). Later, some CMT2A alleles were associated with excessive fusion in patient fibroblasts (Chevrollier et al., 2012, Codron et al., 2016), HeLa cells (Das et al., 2022a, Das et al., 2022b, Das et al., 2024) and fly models (El Fissi et al., 2018). Consistent with our drosophila data, human *MFN2^D2l4N^* and *MFN2^W740S^* were reported as fusion competent (Detmer and Chan, 2007, Li et al., 2019) whereas human *MFN2^K357T^* and *MFN2^R364W^* were associated with enlarged mitochondria in mouse and human cells respectively (Stavropoulos et al., 2021, Das et al., 2022a, Das et al., 2024). The mechanism by which Bulb mutants lead to increased mitochondrial size is unknown and may differ between alleles as they target different protein regions: HB1-HB2 interface for *dD214N*, GD-HB1 and HB1-HB2 interfaces for *dR364W*, hinge 2 for *K357N* and HR2 surface for *W740S*. Recent studies suggested that complex mechanisms could be involved, the hyperfusion caused by *MFN2^R364W^* in human cells being related to an inhibition of mitochondrial fission consecutive to the imbalance between MFN2 and DRP1 ubiquitination (Das et al., 2022a, Das et al., 2022b).

Mfns from yeast, flies and mammals are targets of E3 Ubiquitin ligases regulating Mfn steady state level and functions, in particular mitophagy (Escobar-Henriques and Joaquim, 2019, Alsayyah et al., 2020). In drosophila, Parkin ubiquitinates Marf and negatively regulates its expression level (Ziviani et al., 2010). When overexpressed in CMT2A flies, Parkin attenuated Bulb phenotype but did not counteract mitochondrial clustering in Grape and Tangle. In Tangle mutants, mitochondria became more fragmented showing that wild-type Marf molecules were efficiently targeted by Parkin. Regarding Peanut group, Parkin induced mitochondrial clustering in *dR250Q*. This suggests that degrading wild-type Marf has revealed a cryptic aggregative property, an observation consistent with the semi-dominance described in R250Q patients (Tomaselli et al., 2016). The fact that Parkin is active on Bulb mutants that restores mitochondrial tubules formation in *Marf ^KO^* flies, but not on Grape, Tangle and Peanut alleles that are fusion-defective, suggests a relationship between Marf activity and Parkin-mediated ubiquitination. Interestingly, Fzo1 ubiquitination and subsequent proteasome-dependent degradation are required for mitochondrial fusion in yeast. GD mutations, including a CMT2A-related one, abolish Fzo1 ubiquitination and degradation by impairing Fzo1 interaction with SCF^Mdm30^ ubiquitin ligase (Amiott et al., 2009, Anton et al., 2011, Cohen et al., 2011). Indeed, altered GTPase activity would prevent a conformational change required to reveal an otherwise hidden SCF^Mdm30^ binding site (Cohen et al., 2011). Interestingly, *mdm30* inactivation trigger Fzo1 accumulation, paradoxically associated with mitochondrial fragmentation and aggregation (Cohen et al., 2011). Because promoting Fzo1 degradation independently of SCF^Mdm30^ rescues this phenotype, it was proposed that Fzo1 turnover positively regulates mitochondrial fusion (Cohen et al., 2011). A similar mechanism could contribute to mitochondrial clustering in Grape and Tangle mutants. Consistently, when we looked at the ubiquitination status of mutant Marf by western blot, we found that ubiquitination was reduced in all loss-of-function mutants (*dRl04W*, *dHl28R*, *dL92P*, *dR94W*, *dL248V*, *dH361Y*, *dL745P*). Although the overall decrease was lower, we also found reduced ubiquitination in the mutants that restored mitochondrial tubulation in *Marf ^KO^* (*dR250Q*, *dK357N*, *dR364W* and *dW740S* but not *dD214N*). This suggests that *dR250Q*, *dK357N*, *dR364W* and *dW740S* mutations somehow perturb the functional relationship between Marf and Parkin when Parkin is expressed at steady state level, this alteration being however compensated by Parkin overexpression. Consistent with a role of ubiquitination in Marf turnover, we observed increased expression level for most CMT2A proteins. This observation suggests that defective ubiquitination increases mutant to wild-type protein ratio, possibly contributing to CMT2A alleles dominant properties. However, although the most neurotoxic alleles were the most overexpressed (*dRl04W*, *dHl28R*, *dL92P*), phenotype severity did not correlate with expression levels for the all mutants. This demonstrates that the allele toxicity is primarily determined by its impact on Mfn activity, the increased expression level acting as an amplifier.

We observed varying degrees of neurotoxicity in our allelic series. Consistent with their phenotypes, Grape alleles showed the highest toxicity in flies and Peanut the lowest. Tangle and Bulb mutants had intermediate toxicity which degree varied depending on the allele. Overall, we found that reduced locomotor performance correlated to the extent to which mitochondria had a focal distribution in neuronal cytoplasm. The improper mitochondrial repartition could be due to aggregation or unbalanced fusion depending on mutant group. This mitochondrial sequestration would presumably create dysfunctional cytoplasmic regions by affecting the delivery of mitochondria to the areas where their activity is needed, in particular synapses, and by impairing the functional wiring of mitochondria to the rest of the organelle community. In addition, the abnormal Grape, Tangle and Bulb mitochondria were poorly connected to the network, possibly exacerbating the effects of their inefficient distribution. Importantly, phenotype severity in flies correlated well with disease severity in human. Grape mutants (*dR94W*, *dRl04W*, *dHl28R*, *dL248V*) as well as Tangle and Bulb mutants (*dL92P*, *dH36lY*, *dR364W*, *dK357N*) with the lower locomotion in flies corresponds to MFN2 mutations causing severe and early onset CMT2A (Kijima et al., 2005, Calvo et al., 2009, Stuppia et al., 2015). Conversely, the less affected Tangle (*dR259L*) and Bulb (*dD214N*, *dW740S*) mutants as well as Peanut mutants (*dR250Q*, *dL745P*) correspond to MFN2 mutations associated with mild and late onset CMT2A (Nicholson et al., 2008, Ajroud-Driss et al., 2009, Nakhro, 2013, Stuppia et al., 2015). Our data support the pathogenicity of four human alleles whose involvement in CMT2A was classified as uncertain (D214N, R250Q, R400P, and L745P), as the related mutations in flies affect MARF activity. Regarding L745P, the fly-related mutation causes a loss of function without dominant properties. This suggests a sensitizing condition that might trigger haploinsufficiency in patients. Finally, the phenotypes of *dR250Q* and *dD214N* flies very well echo the clinical data. Indeed, the R250Q and D214N alleles only trigger CMT2A when patients are compound heterozygous and were proposed to act as semi-dominant (Nicholson et al., 2008, Piscosquito et al., 2015, Tomaselli et al., 2016).

Taken together, our findings support the use of drosophila as a test tube to validate the implication of novel MFN2 genetic variants in CMT2A and predict their putative severity. Overall, these data show a strong correlation between phenotype strength in flies and disease severity in humans, supporting the relevance of drosophila in the study of CMT2A disease.

## Methods

### Protein sequence analysis and structure modelling

BLAST was used to retrieve the sequences of Mfns (Marf: 11 insect species ; MFN1 and MFN2: 11 vertebrate species) and of BDLPs (26 *Nostoc* species). BDLPs were aligned using T-Coffee (Notredame et al., 2000) simple alignment tool based on ClustalW algorithm. Mfns were aligned using the PSI-Coffee (Notredame et al., 2000). Aa identity and similarity were calculated with SMS Ident and Sim (https://www.bioinformatics.org/sms2/ident_sim.html) using default parameters (GAVLI, FYW, CM, ST, KRH, DENQ, P). Outputs of these two alignments were merged using the T-Coffee Combine alignment algorithm. MFN1, MFN2, Marf and BDLP linear alignment was manually refined according to the structural alignment of MFN2 (PDB: 6JFK) and BDLP (PDB: 2J68) (Low and Lowe, 2006, Li et al., 2019) obtained using ChimeraX (Goddard et al., 2018). This alignment was used for template-based molecular modelling of MFN2 and Marf using BDLP structures in open and closed conformations (PDB: 2J68, 2W6D) (Low and Lowe, 2006, Low et al., 2009). As MFN2 and Marf have longer N-terminal tails than BLDP, we built N-terminal truncated models: MFN2-Δ1-23 and Marf-Δ1-60, respectively. Transmembrane regions were predicted with TMpred (Cuthbertson et al., 2005). A single-template modelling script was used to generate over 100 of closed-and open-conformation models using the MODELLER program (v 10.0) (Sali and Blundell, 1993). The GDP ligand was docked to the catalytic pocket either by modelling procedure, or using ChimeraX structure analysis tool (https://www.cgl.ucsf.edu/chimerax/). In the end, one homology model was selected based on the lowest *modpdb* value for the closed and open conformations of MFN2 and Marf. The hydrogen were then added on a MolProbity (Chen et al., 2010). The stereochemical quality was assessed using MolProbity and ProCheck. Z-score and local energy quality were determined using ProSA (Wiederstein and Sippl, 2007). Models of Marf dimers were generated using machine learning-based modelling (Jumper et al., 2021) with AlphaFold3 algorithm (https://alphafoldserver.com/). The resulting model was in a GDP-bound-like state (GD tilted upward) as found in resolved GDP-bound MFN2 and MFN1 truncated structures (Yan et al., 2018, Li et al., 2019). Marf with GTP-bound-like state (GD tilted downward) was generated with AlphaFold2, providing truncated MFN1 GTP-bound structure (PDB: 5YEW) as a template and reducing multiple alignment size (*max_msa*) (setting: 16:32) (https://colab.research.google.com/github/sokrypton/ColabFold/blob/main/AlphaFold2.ipyn). Graphical representation and residue mapping were generated using ChimeraX.

### *Drosophila* stocks and culture

#### *Marf UAS* lines

*Marf* coding sequence was amplified from *w^lll8^* fly cDNAs and inserted into the pUASt-derived vector pPHW (from T. Murphy) by Gateway cloning using pDonr221 as intermediate. This *Marf* sequence differed from the one on Flybase by silent mutations (natural polymorphisms and a synthetic EcoNI site introduced for screening convenience) (Supplementary sequence). *Marf* 5’ and 3’ UTRs were not included. A CACC kozak sequence (Acevedo et al., 2018) and three N-terminal HA tags were added. All CMT2A mutations were made by PCR (CloneAmp HIFI, Takara Bio). Constructs were verified by Sanger sequencing. Vectors carrying *Marf* wild-type or mutant alleles were inserted into chromosome 2L (25C6) by phiC31-mediated transgenesis using *w* ; *attP40* flies (Bestgene). Strains from BDSC: *elav-GAL4* (#458), *OK371-GAL4* (#26160), *UAS-myr::mRFP* (#7119), *UAS-KDEL::GFP* (#9898), *UAS-parkin_IR* (#38333, TRiP.HMS01800) and Y *3xP3-DsRed2* (#41459). The following lines were previously described by the authors: *UAS-mit::dendra2*, *Marf KO* (El Fissi et al., 2018) and *UAS-mit::mKate2* (Poliacikova et al., 2023). *UAS-Parkin* was a generous gift from Ming Guo. GAL4/UAS fly crosses were grown at 29°C on a cornmeal-agar diet. We used third instar wandering larvae, except for *Marf ^KO^* larvae which stage could not be precisely determined as Mfn loss of function alters larval developmental (Sandoval et al., 2014, El Fissi et al., 2018).

### Adult fly emergence quantification

#### elav-GAL4 I Y

*UAS-mit::dendra2* males were crossed to *UAS-Marf^CMT2A^ ^or^ ^WT^* females providing a progeny in which females but not males expressed CMT2A alleles as *elav-GAL4* is inserted on the X chromosome. Developmental lethality was assessed by counting the number of females and males in the adult offspring. Similar approach was applied to crosses made using *OK37l-GAL4ICyO* flies as only the wild-type wing progeny express GAL4.

### Analysis of photoreceptor degeneration

#### elav-GAL4

*UAS-Marf^CMT2A^ ^or^ ^WT^ I UAS-mit::dendra2* female flies were anesthetised with CO_2_ and decapitated with a micro-scissor. The back of the head was immediately glued with methyl cyanoacrylate onto a glass slide and the eyes covered with immersion oil (Leica Type F). Observations were made on a Zeiss Axioplan 2 microscope at 63X with light at maximum intensity and diaphragm at lower aperture. Photoreceptor degeneration was scored by counting the number of ommatidia with less than seven rhabdomeres. We analysed on average 175 ommatidia from 8 individuals per condition. Statistical analysis was performed using Mann-Whitney U test.

### Locomotor assay

Fly development was carried out at 29°C, except for *dR94W*, *dR104W*, and *dH128R* that were grown at 25°C to reduce developmental lethality and get adults. Adult flies (*elav-GAL4 ; UAS-Marf^CMT2A^ ^or^ ^WT^ I UAS-mit::dendra2*) were aged 10 days by group of 15 flies, the culture vials being changed every two days. Locomotor performance was determined by startle-induced negative geotaxis (SING). We used an empty culture vial in which one piece of PU foam plug was inserted at the bottom and one at the top defining a free space of 65 mm height. A dozen of flies were introduced in the vial, allowed to recover from CO_2_ anaesthesia (30 min) and tapped down to the bottom to trigger SING. A picture of the tube was taken 5 sec later using a digital camera. The height reached by each flies was measured using ImageJ. To test for the effect of vision on SING, a group of flies was first assessed into the complete darkness and then in the light. Three to four independent races involving a total of 20-30 flies were performed per condition. Statistical analysis was performed using Mann-Whitney U test.

### Live confocal imaging

Third instar larvae tissues were very rapidly dissected, mounted and observed under the microscope (<5 min). *Marf ^KO^* males were sorted thanks to a *3xP3-mCherry* Y chromosome. CNS or ventral cuticle were mounted in PBS. Two bands of double-face tape were placed on both side of the samples to avoid crushing between the coverslips and the slide. We imaged dorsal motor neuron clusters of the ventral nerve cords and NMJs located at the surface of muscles attached to the ventral cuticle. Z-image stacks (interval 0.2 µm) covering the whole depth of motor neuron clusters or NMJs were acquired under a Zeiss LSM880 confocal microscope at low laser intensity (<10%) with 63X (NMJs) or 100X (cell bodies) objectives. For high resolution observations, we used a Airyscan detector.

### Mitochondrial morphology analysis

Airyscan-acquired Z-image stacks of living motor neurons (dorsal clusters of larval ventral nerve cord) expressing mit::dendra2 were submitted to deconvolution (Zen 2.3SP1 software with auto-settings). To analyse mitochondria in a wild-type background, we used Z-stacks, Z-projection (Fiji, Temporal Colour Coder) and 3D reconstruction (Fiji, 3D project) as templates for scoring in blind pre-defined morphological mitochondrial traits i.e. fragmented, filamentous, enlarged, net-like and clustered. To analyse mitochondria in Peanut and in *Marf ^KO^* flies, mitochondria were automatically segmented on confocal Z-projections of 10 images using a Weka classifier (Fiji, https://imagej.net/plugins/tws/). Then, total branch length and number were determined using MitoAnalyser (Fiji, https://github.com/AhsenChaudhry/Mitochondria-Analyzer) with default parameters.

### Hierarchical clustering

First, we submitted to hierarchical clustering the quantifications of mitochondrial shape obtained when CMT2A mutations were expressed in a wild-type background. In these quantifications, the presence of 5 mitochondrial traits was scored in each cell analysed (see above). Using the R package *stats* version 4.1.3, the distance matrix between each cell was calculated using the *dist* function with the *binary* method (as data were represented by the absence or presence of a mitochondrial trait). Then, a mean distance was calculated between each mutation. Then, the distance matrix between each mutation in a *Marf ^KO^* background was calculated for the mean mitochondrial length, for the percentage of fragmented mitochondria (length below 0.6 µm) and for the percentage of filamentous mitochondria (length above 1.4 µm) using the *dist* function with the *euclidean* method. Then, a hierarchical clustering between this new matrix and the previous matrix of mean distance of the presence/absence of the five mitochondrial traits was calculated using *hclustcompro* with the method *ward.D* from the R package *SPARTAAS* version 1.2.4 (https://CRAN.R-project.org/package=SPARTAAS), which is a tool allowing to combine distance matrices from two different sources to create this clustering and defined potential clusters. We let the tool calculates the α parameter, which is the mixing parameter giving different weights to each matrix (here α=0.59, giving more weight to the wild-type background matrix). Finally, the dendrogram created was also converted to newick format using the package *ape* version 5.7-1 (Paradis and Schliep, 2019) and visualised using SeaView version 5.0.5 (Gouy et al., 2021).

### Electron Microscopy

Larval CNS were rapidly dissected, placed on a glass slide, incubated in a drop of fixation solution (FS) (2% paraformaldehyde, 2.5% glutaraldehyde, 5mM CaCl_2_, 0.1mM Na cacodylate) for 30 min at room temperature and embedded in a drop of 2% low melting agarose. Agarose cubes containing larval CNS were cut out with a scalpel and placed in FS for 24 hr at 4°C. This was followed by 2 hr post-fixation (2.5% glutaraldehyde, 0.8% osmium tetroxide, 0.1mM Na cacodylate) at 4°C. Ultrathin Epon plastic sections were contrasted in 2% uranyl acetate for 10 min and lead citrate for 2.5 min and observed with a Tecnai G2 TEM at 200 kV.

### Mitochondrial content mixing

Dissected CNS from *OK371-GAL4, 2 X [UAS-mit::dendra2 IUAS-HA::Marf]* larvae were observed under a Ropper-Nikon spinning disc confocal microscope equipped with photo-conversion module controlled by Metamorph. Mit::dendra2 was photoconverted at 450nm with 5% laser power for 1800 ms (300 iterations). Green (excitation 488nm) and red (excitation 545nm) mit::dendra2 signals were acquired repetitively during a time window of 180 sec. The photoconverted areas of the first image (area Ai) and last image (area Af) were measured using Fiji to calculate a diffusion index Af/Ai expressed as percent of control.

### Mitochondrial and ER distribution

Confocal images of *OK371-GAL4, UAS-mit::dendra2IUAS-HA::Marf ; UAS-myr::mRFP* third instar larvae were cropped around individual motor neurons according the myr::mRFP signal and the size of all images adjusted to 128×128 pixel. The mit::dendra2 signal was automatically thresholded (Otsu method), and all pixel *x* and *y* coordinates were extracted. The variance of x and y coordinates was calculated and summed (*x*+*y* variance). For each condition, an average of nine cell bodies were analysed and the mean *x*+*y* variance was calculated and expressed as percent of control to obtain a mitochondrial distribution index. The same approach was used for ER distribution based on KDEL::GFP signal.

### Manders’ correlation between ER and mitochondria staining

Confocal images of motor neuron of *OK371-GAL4, UAS-KDEL::GFP,UAS-mit::mKate2IUAS-HA::Marf* third instar larvae were automatically treated to remove background on each channel (Fiji, rolling ball method). Manders’ correlation coefficient between mit::mKate2 (mitochondria) and KDEL::GFP (ER) signals was then determined (Fiji, Coloc2 plugin).

### NMJ mitochondrial content

NMJ Z-stacks were treated using Fiji to create Z-projections (maximum intensity) and perform an automatic threshold of the mit::dendra2 and myr::mRFP channels (Otsu method). The integrated pixel density, that is an expression of the area on binary images, was calculated for each bouton in each of the channel. The integrated density ratio, corresponding to the ratio of the mitochondria area over the bouton area, was calculated. On average a hundred NMJ boutons were analysed in each condition.

### Correlograms

Using the *cor* function from the R package stats version 4.1.3, we calculated the Kendall correlation coefficient between each quantitative variable. We chose the Kendall correlation coefficient over Pearson or Spearman because it is not parametric and avoid any assumptions about the data distribution. Then, these coefficients were plotted using the *corrplot* function from the R package *corrplot version 0.92* (https://github.com/taiyun/corrplot).

### Code availability

The code used to generate the results is available in our Github repository (https://github.com/cedricmaurange/Varuk-et-al.git)

### Antibodies for western blots

Primary antibodies: rabbit anti-Marf (gift from Alex Whitworth) (Ziviani et al., 2010) (1/1,000), rat anti-HA (Roche, clone 3F10) (1/1,000), rat anti-α-Tubulin (Abcam, Ab6160) (1/10,000), mouse anti-Ubiquitin (Merck, clone FK2) (1/500). Secondary HRP-coupled antibodies: rabbit anti-rat (Jackson, 312-005-003), donkey anti-mouse (Jackson, 715-036-151), mouse anti-rabbit (Jackson, 211-032-171) (all at 1/20,000).

### CNS protein extraction

Five third instar larva CNS were dissected, immediately immersed into 20 µL of Laemmli buffer containing 1mM DTT and protease inhibitors (Complete, Roche), heated 20 min at 70°C, vigorously vortexed and stored at -20°C. Prior loading, protein samples were heated at 95°C for 5 min.

### Immunoprecipitation

Twenty five third instar larva CNS expressing HA::Marf were immersed in 100 µL of cold lysis buffer (PBS, 0.1% NP40, Complete protease inhibitor from Roche and 1mM DTT) and frozen in liquid nitrogen. The operation was repeated three times to obtain 75 CNS. The samples were rapidly thawed at 37°C and snap frozen in liquid nitrogen three times, and then pooled and homogenised using a 1 mL glass grinder and a tight Teflon pestle. The resulting suspension was centrifuged twice at 800 g and the supernatant frozen at -80°C before immunoprecipitation (IP). For IP, the sample was incubated 2 h at 4°C with rat anti-HA antibody (Roche, clone 3F10) (1/40). Protein G-coupled magnetic beads (Dynabeads, Thermo Fisher) were incubated for 45 min at 4°C with rabbit anti-rat antibody (Jackson, 312-005-003) (1/20) and then mixed with the HA-conjugated protein sample for 30 min at 4°C. The beads were washed according to manufacturer instructions and immunoprecipitated HA::Marf was recovered following a 10 min incubation at 95°C in 6X Laemmli sample buffer containing 5% of freshly added β-mercaptoethanol.

### Western Blotting

Proteins were separated on a 10% polyacrylamide gel in the presence of SDS at 100V. Transfer was done at 100V for 1.5 h either on nitrocellulose membrane for larval CNS samples, or on ethanol activated PVDF (0.45 µm) membrane for IP samples in 25mM tris-base, 190mM glycine supplemented with ethanol (20% for nitrocellulose, 10% for PVDF). For anti-Ubiquitin staining on IP samples, PVDF membranes were boiled after transfer for 10 min in distilled water (Emmerich and Cohen, 2015). Blots were saturated in 5% BSA in PBS Tween20 0.05% (PBT), probed with primary antibody and corresponding HRP-coupled secondary antibodies with washing steps performed in PBT. Signal was revealed with chemiluminescent substrate (SuperSignal West Pico Plus for anti-HA and Atto for anti-Ubiquitin, Thermo Fisher) and images were acquired with a Bio-Rad imager. Blots of IP samples labelled with anti-Ubiquitin were stripped 7 min in Restore WB Stripping Buffer (Thermo Scientific), washed and relabelled with anti-HA. Band pixel intensities were measured using ImageJ (gel analysis plugin) and quantification were performed on at least three independent blots with independent samples. α-Tubulin was used for normalisation.

## Acknowledgements

We thank Jean-Pierre Duneau (LISM, Marseille) for sharing his expertise on protein structure modelling and the IBDM microscopy and fly facilities for technical support. The IBDM-PiCsL platform is supported by ANR for the future program (France-BioImaging, ANR-10-877 INSB-04-01). We are grateful to Alice Corbet, Elisa Montredon and Carole Rubio for technical involvement. Experimental expenses, OOV’s PhD fellowship, AR and CR’s salaries, and CO and AC’s internships were founded by the Association Fran;:aise contre les Myopathies (AFM-Telethon) (grant 22241 and 20097 to TR and fellowship 22070 and 23855 to OOV). We thank Lydia Kerkerian-Le Goff for scientific advices, and Laurent Kodjabachian and Pascale Durbec for supporting OOV’s PhD completion with IBDM funding. Stocks from the Bloomington Drosophila Stock Center were used (NIH P40OD018537).

## Author contribution

Project conception, supervision: TR. Experiment design: OOV, TR. Confocal, EM imaging: OOV, TR. Image analysis: OOV, TR, AC, AR. Lethality, locomotion, neurodegeneration: CO, AC. Biochemistry: OOV, TR. Fly genetics: OOV, AR, TR. Molecular biology: AR, OOV. EM samples: AA. Clustering, correlation: PdB. Lab logistics: AR, OOV. Manuscript: TR, OOV.

