## Supplementary Material for "Multifaceted fusion defects converge on altered mitochondrial distribution and increased mutant mitofusin level in CMT2A models"

### Supplementary Data

**Figure S1. Secondary structures targeted by CMT2A mutation hotspots.**

**(A)** Frequency of CMT2A mutation along the aa sequence of human MFN2 showing hotspots of mutations in specific secondary structures. Heat map correspond to the proportion of mutated aa within a specific secondary structure. **(B)** Resolved structure of human MFN1 dimers (Yan et al., 2018) in the GTP-bound/open conformation (PDB: 5YEW). To superimpose CMT2A mutation heat map shown Figure 1 on MFN1 structures, MFN1 and MFN2 sequences were aligned and the frequency of mutations for a given 10 aa interval of MFN2 was assigned to the homologous interval in MFN1. Arrows indicate the most affected secondary structures. **(C)** Resolved structure of human MFN1 dimers (Yan et al., 2018) and the GDP-bound/closed conformation (PDB: 5GOM) (bottom panel). MFN2 CMT2A mutation heat map was superimposed based on sequence alignment. **(D)** Resolved structure of BDLP dimers in open conformation inserted in lipid bilayer (Low and Lowe, 2006) (PDB: 2W6D). MFN2 CMT2A mutation heat map was superimposed based on sequence and structural alignments between MFN2 and BDLP (see methods). **(E)** Resolved structure of BDLP dimers in closed conformation (Low and Lowe, 2006) (PDB: 2W6D). MFN2 CMT2A mutation heat map was superimposed based on sequence and structural alignments between MFN2 and BDLP (see methods).

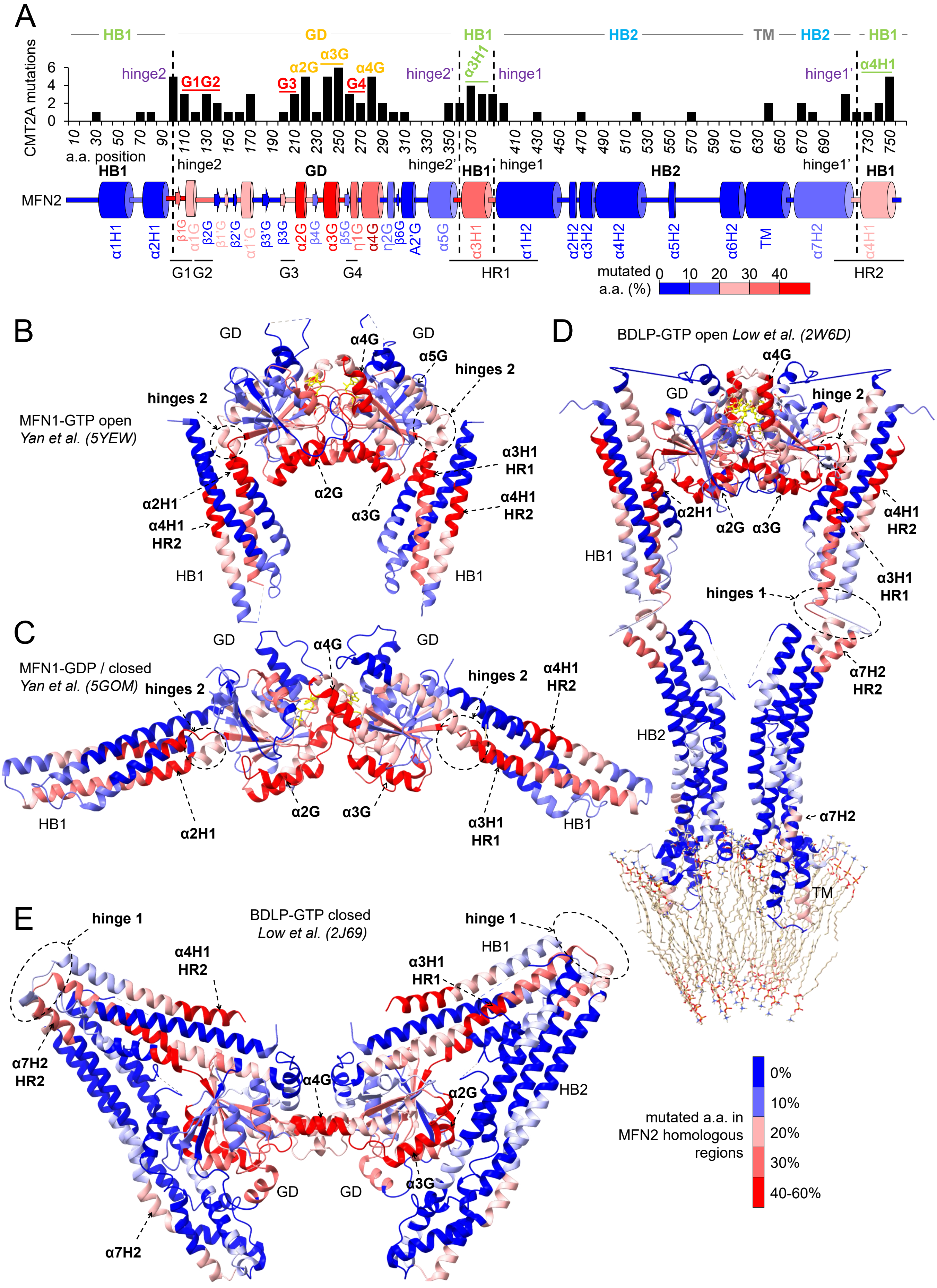
A
HB1
GD
HB1
HB2
TM
HB2
HB1
CMT2A mutations
hinge2
G1G2
G3
α2G
α3G
G4
α4G
hinge2'
α3H1
hinge1
hinge1'
α4H1
a.a. position
hinge2
GD
hinge2'
HB1
hinge1
HB2
hinge1'
HB1
MFN2
G1G2
G3
G4
HR1
HR2
mutated a.a. (%)
B
MFN1-GTP open
Yan et al. (5YEW)
hinges 2
α4G
α5G
α2G
α3G
α3H1 HR1
α4H1 HR2
α2H1
HB1
C
MFN1-GDP / closed
Yan et al. (5GOM)
D
BDLP-GTP open Low et al. (2W6D)
hinge 2
hinges 1
α7H2 HR2
HB2
TM
E
hinge 1
BDLP-GTP closed
Low et al. (2J69)
α7H2
mutated a.a. in MFN2 homologous regions
0%
10%
20%
30%
40-60%

**Figure S1**

**Figure S2. Most amino acids mutated in CMT2A are conserved in Marf.**
**(A)** Alignment of aa sequences from human MFN1 and MFN2 (UniProt: Q8IWA4 and O95140) (hMFN1 and hMFN2) and *D. melanogaster* Marf (UniPrto: Q7YU24) (dMarf). Black: identical residues between Marf and/or MFN1 and MFN2 sequences. Dark pink in MFN2: residues mutated in CMT2A. Dark pink in Marf: residues identical to CMT2A-involved aa. Pale pink in Marf: residues with physicochemical properties similar to CMT2A-involved ones. Red frames: G1, G2, G3, G4 motifs. In G2, MFN2-T129, Marf-T170 and MFN1-I108 are highlighted in blue. Black frames: HR1 and HR2 motifs. HB1 and GD α-helices are shown as cylinders and GD β-strands as arrows following MFN2 truncated structure (PDB: 6JFK) (Li et al., 2019). HB2 α helices are predicted from BDLP and TM from TMpred. Numbers corresponds to the positions of CMT2A aa substitutions, the ones that have been coloured being selected for investigation in drosophila. **(B)** Percentage of identity and similarity between the sequences from drosophila Marf (Q7YU24), human MFN2 (O95140), considering all aa (hMFN2) or only those mutated in CMT2A (hMFN2-CMT2A), and human MFN1 (Q8IWA4) (hMFN1). **(C)** Positions of CMT2A-related residues selected for our study in flies in the sequence of human MFN2, drosophila Marf and human MFN1 based on sequence alignment.

A

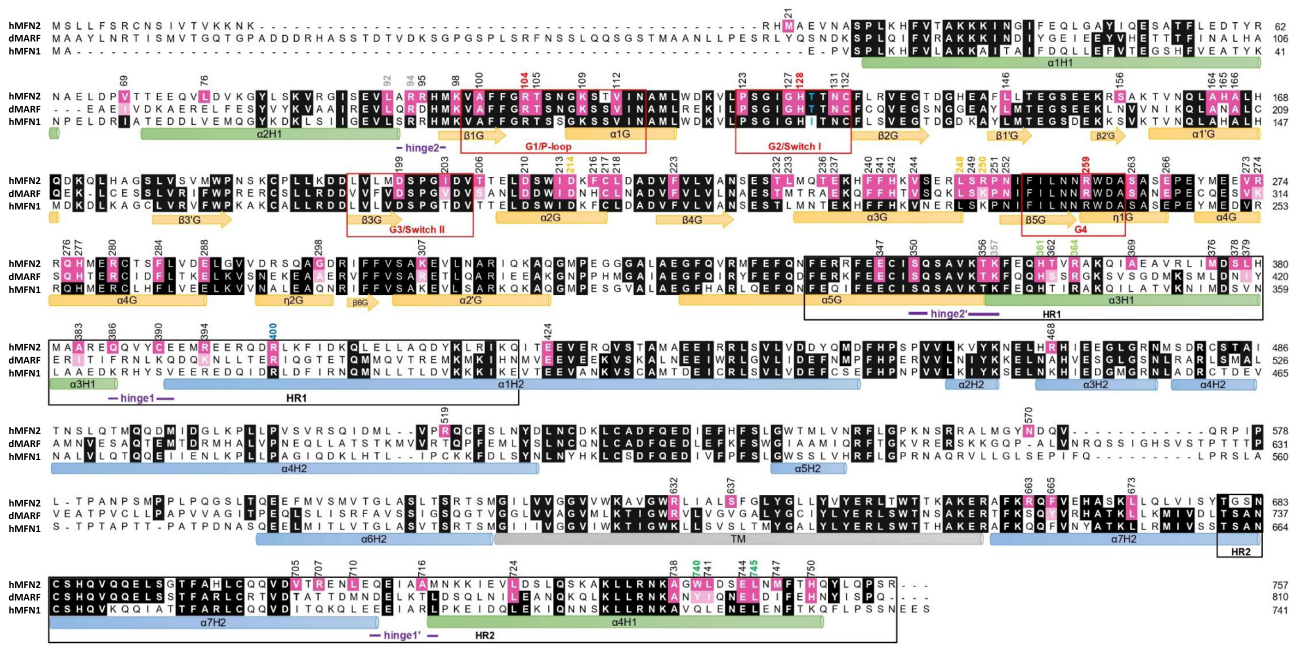

B

| | Identity (%) | Similarity (%) | Total (%) |
|---|---|---|---|
| **dMarf / hMFN2** | **45.52** | **15.22** | **60.74** |
| **dMarf / hMFN2-CMT2A aa** | **60.64** | **13.82** | **74.46** |
| dMarf / hMFN1 | 42.37 | 13.67 | 56.04 |
| hMFN2 / hMFN1 | 58.51 | 13.61 | 72.12 |

C

| | | | | | | | | | | | | | | |
|---|---|---|---|---|---|---|---|---|---|---|---|---|---|---|
| **hMFN2** | L92 | R94 | R104 | H128 | D214 | L248 | R250 | R259 | K357 | H361 | R364 | R400 | W740 | L745 |
| **dMarf** | L133 | R135 | R145 | H169 | D254 | L288 | K290 | R299 | K397 | H401 | R404 | R440 | Y794 | L799 |
| **hMFN1** | L71 | R73 | R83 | H107 | D193 | L227 | K224 | R238 | K336 | K340 | R343 | R379 | Q721 | L726 |

***GD*** ***Cat. Site*** ***HB1*** ***Hinge*** ***HB2***

**Figure S2**

**Figure S3. Models of Marf protein structures**

**(A)** Marf model (aa 1- 60 deleted) built by homology modelling based on BDLP GTP-bound (GMPPNP ligand) open conformation. Spheres correspond to the lateral chain atoms (except hydrogens) of aa that have been mutated in flies to mimic CMT2A alleles. **(B)** Marf model (aa 1- 60 deleted) based on BDLP GDP-bound closed conformation. Spheres correspond to the lateral chain atoms (except hydrogens) of aa that have been mutated in flies to mimic CMT2A alleles. **(C)** Marf model based on BDLP GTP-bound (GMPPNP ligand) open conformation. Highlight on GD and HB1 interface (side view). Sticks: aa lateral chains. **(D)** Marf BDLP-based model. Highlight on GD catalytic site (front view). Sticks: aa lateral chains. **(E)** Marf BDLP-based model. Highlight on HB1 (top view). Sticks: aa lateral chains. **(F)** Model of Marf dimer structure (aa 1- 60 deleted) predicted by machine learning-based modelling (AlphaFold3). Spheres: lateral chain atoms (except hydrogens) of aa that have been mutated in flies to mimic CMT2A alleles. Top panel: side view. Bottom panel: top view.

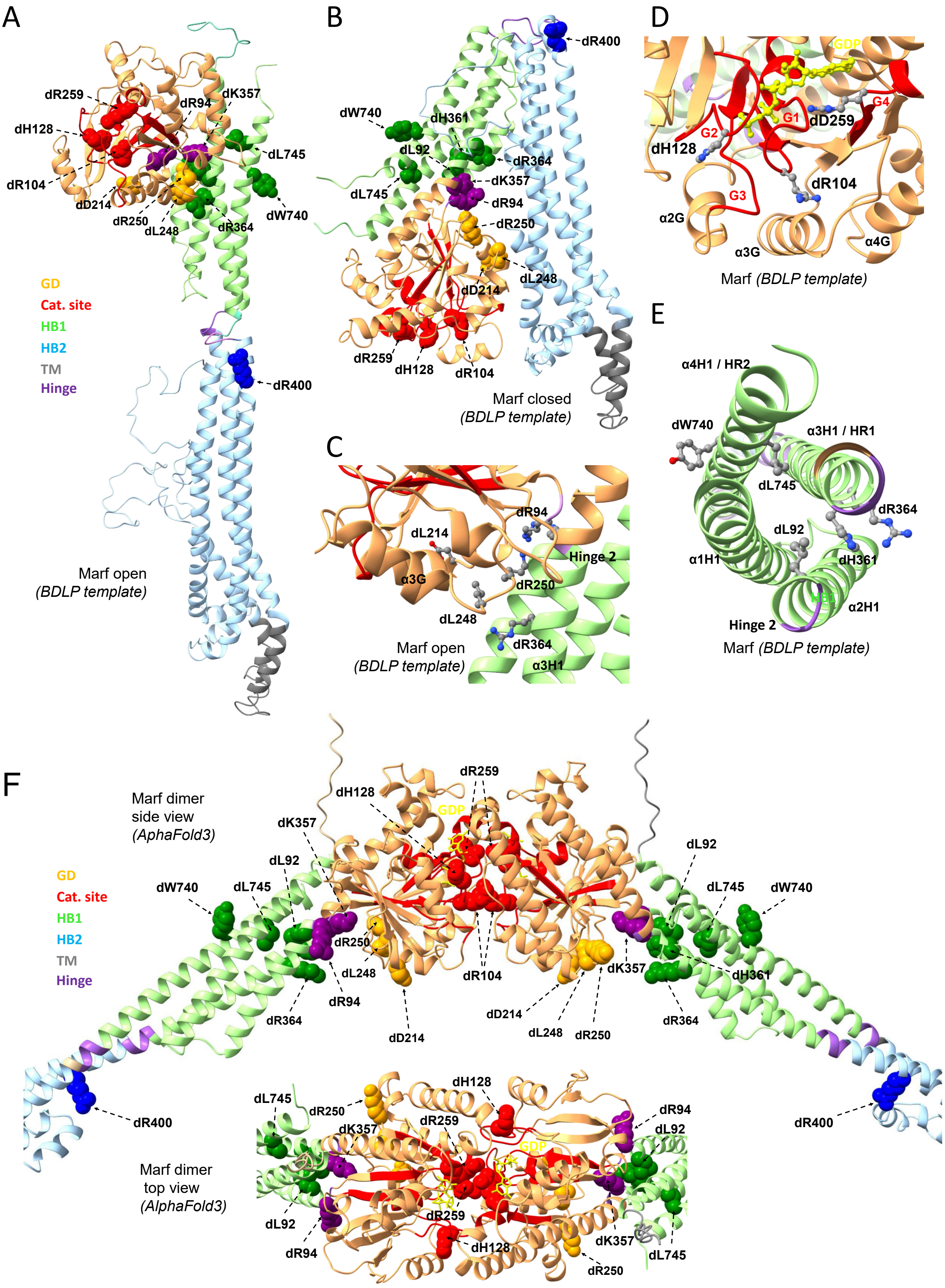
A
dR259
dH128
dR104
dD214
dR250
dL248
dR94
dK357
dL745
dW740
dR364
GD
Cat. site
HB1
HB2
TM
Hinge
dR400
Marf open
(BDLP template)
B
dR400
dW740
dH361
dL92
dL745
dR364
dK357
dR94
dR250
dL248
dD214
dR259
dH128
dR104
Marf closed
(BDLP template)
C
dR94
dL214
α3G
Hinge 2
dR250
dL248
dR364
α3H1
Marf open
(BDLP template)
D
GDP
G4
G1
dD259
G2
dH128
G3
dR104
α2G
α3G
α4G
Marf (BDLP template)
E
α4H1 / HR2
dW740
α3H1 / HR1
dL745
dR364
dL92
α1H1
dH361
α2H1
Hinge 2
Marf (BDLP template)
F
Marf dimer
side view
(AphaFold3)
GD
Cat. site
HB1
HB2
TM
Hinge
dR259
dH128
GDP
dK357
dL92
dW740
dL745
dR250
dL248
dR94
dR364
dD214
dR104
dK357
dH361
dD214
dL248
dR250
dR364
dL92
dL745
dW740
dR400
dR400
Marf dimer
top view
(AlphaFold3)
dH128
dL745
dR250
dK357
dR259
GDP
dR94
dL92
dL92
dR94
dR259
dH128
dK357
dL745
dR250

### Figure S3

**Figure S4. Comparison of CMT2A-related residues lateral chain localisation and orientation in Marf models and resolved structures of MFN2 and MFN1.**
**(A)** Front view of the catalytic site in AlphaFold3 Marf model and resolved structure of MFN2. **(B)** Top view of the dimer interface formed by GD in AlphaFold3 Marf model and resolved structure of MFN1. **(C)** Side view of GD and HB1 in AlphaFold3 Marf model and resolved structure of MFN2 GDP-bound state. **(D)** Side view of GD and HB1 in AlphaFold3 Marf model generated using an MFN1 template in GTP-bound state (GDP-$BeF_3^-$ ligand) and resolved structure of MFN1 GTP-bound. **(E)** Top view of HB1 in AlphaFold3 Marf model and resolved structure of MFN2 GDP-bound. **(F)** Bottom view of the GD in AlphaFold3 Marf model and resolved structure of MFN2 GDP-bound. **(G)** Colour code.

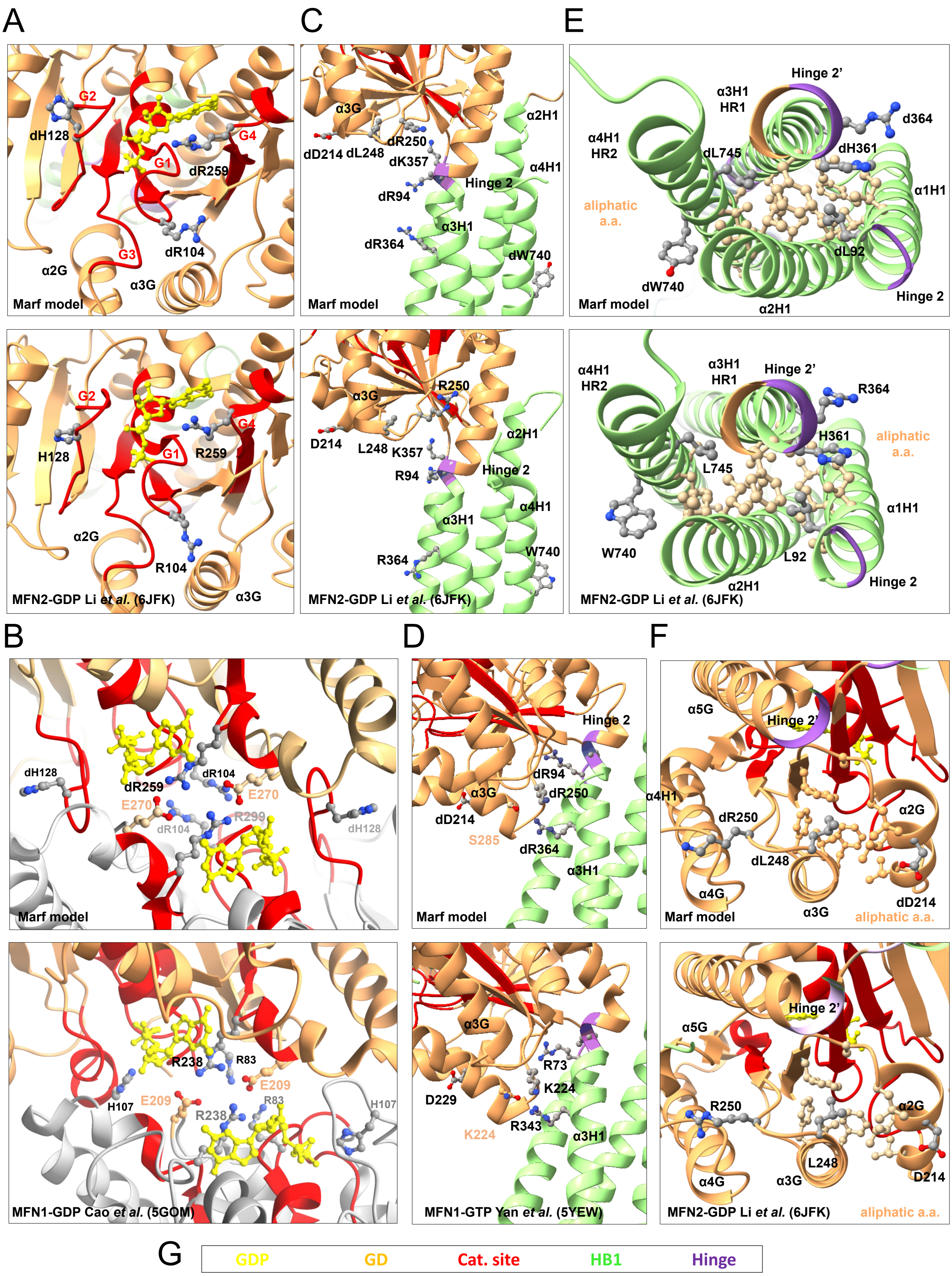
A
G2
dH128
G4
G1
dR259
G3
dR104
α2G
α3G
Marf model
G2
H128
G4
G1
R259
α2G
R104
α3G
MFN2-GDP Li et al. (6JFK)
C
α3G
α2H1
dD214 dL248
dR250
dK357
α4H1
Hinge 2
dR94
α3H1
dR364
dW740
Marf model
R250
α3G
D214
L248
K357
α2H1
Hinge 2
R94
α4H1
α3H1
R364
W740
MFN2-GDP Li et al. (6JFK)
E
Hinge 2'
α3H1
HR1
d364
α4H1
HR2
dL745
dH361
α1H1
aliphatic
a.a.
dL92
dW740
Hinge 2
Marf model
α2H1
α4H1
HR2
α3H1
HR1
Hinge 2'
R364
H361
aliphatic
a.a.
L745
α1H1
W740
L92
α2H1
Hinge 2
MFN2-GDP Li et al. (6JFK)
B
dH128
dR104
dR259
E270
E270
R299
dR104
dH128
Marf model
R238
R83
H107
E209
E209
R238
R83
H107
MFN1-GDP Cao et al. (5GOM)
D
Hinge 2
dR94
α3G
dR250
α4H1
dD214
S285
dR364
α3H1
Marf model
α3G
R73
K224
D229
K224
R343
α3H1
MFN1-GTP Yan et al. (5YEW)
F
α5G
Hinge 2'
α2G
dR250
dL248
α4G
α3G
dD214
Marf model
aliphatic a.a.
Hinge 2'
α5G
R250
α2G
L248
α4G
α3G
D214
MFN2-GDP Li et al. (6JFK)
aliphatic a.a.
G
GDP
GD
Cat. site
HB1
Hinge

**Figure S4**

**Figure S5. Neuronal overexpression of HA-tagged Marf in larval CNS.**
**(A)** Scheme of the UAS/GAL4 system used to express wild-type (*Marf*$^{CTRL}$) and mutant (*Marf*$^{CMT2A}$) forms of HA-tagged Marf in all neurons (*elav-GAL4*) or in motor neurons (*OK371-GAL4*). **(B)** Anti-Marf and anti-α-Tubulin western blot analysis of larval CNS protein extracts from *Marf*$^{KO}$, control flies expressing only endogenous Marf (white arrow) and *elav-GAL4, UAS-HA::Marf*$^{CTRL}$ flies overexpressing HA::Marf (black arrow). **(C)** Fold protein expression of HA-tagged Marf relative to endogenous Marf. Mean of 6 independent samples +/- SD.

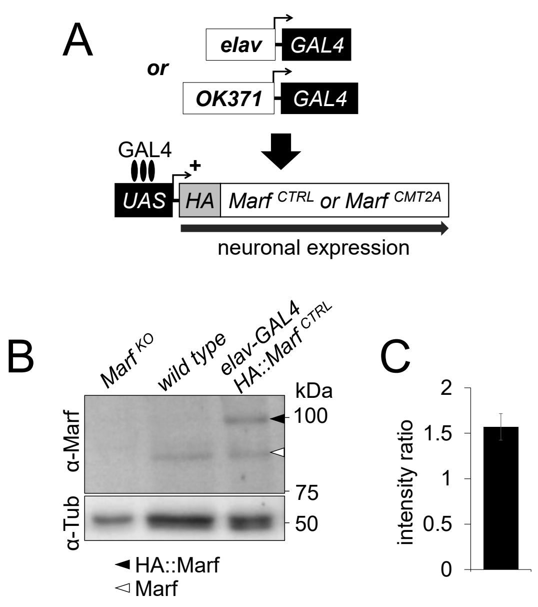
A
elav
GAL4
or
OK371
GAL4
GAL4
+
UAS
HA
Marf CTRL or Marf CMT2A
neuronal expression
B
Marf KO
wild type
elav-GAL4
HA::Marf CTRL
kDa
100
75
50
α-Marf
α-Tub
HA::Marf
Marf
C
intensity ratio
2
1.5
1
0.5
0

**Figure S5**

**Figure S6. Mitochondrial phenotypes associated with CMT2A mutations in a wild-type background.**

Confocal Airyscan image Z-projections (20 slices) of mitochondrial networks (mit::dendra2) from living motor neuron (dorsal clusters of ventral nerve cord) of *OK371-GAL4, UAS-mit::dendra2* third instar larvae. Colour code: Z-depth in dorso-ventral axis. Intermediate panels: high magnification optic slices of mitochondria structures pinpointed by arrows on upper panels. Bottom panels: TEM mitochondria high magnification views in motor neurons (dorsal clusters) from larval ventral nerve cords. **(A)** wild type (endogenous *Marf* only). *Marf* $^{CTRL}$ (*UAS-HA::Marf* $^{WT}$). **(B)** Grape alleles (*UAS-HA::Marf* $^{CMT2A}$). **(C)** Tangle alleles (*UAS-HA::Marf* $^{CMT2A}$). **(D)** Bulb alleles (*UAS-HA::Marf* $^{CMT2A}$). **(E)** Peanut alleles (*UAS-HA::Marf* $^{CMT2A}$).

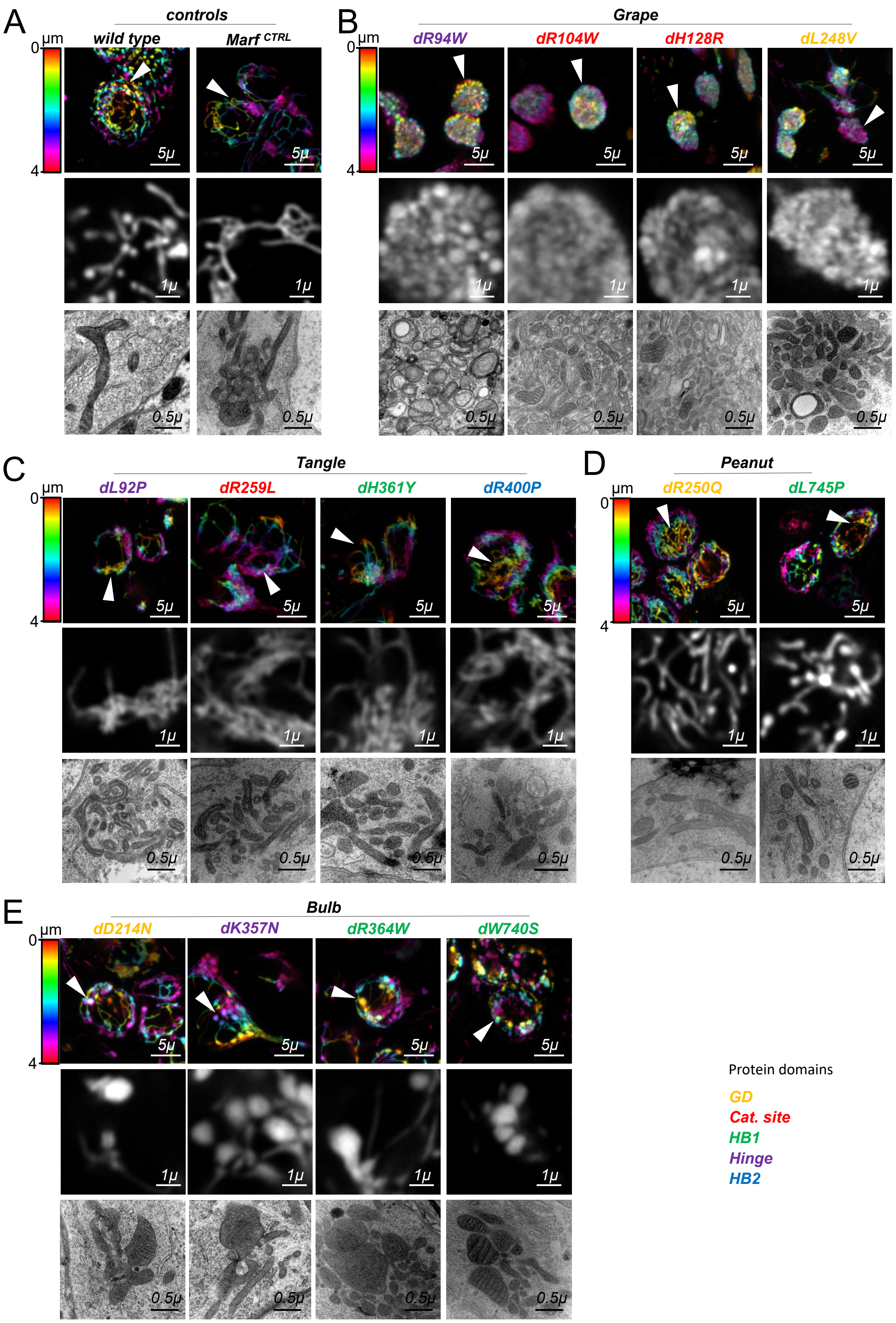
A
controls
wild type
Marf CTRL
B
Grape
dR94W
dR104W
dH128R
dL248V
C
Tangle
dL92P
dR259L
dH361Y
dR400P
D
Peanut
dR250Q
dL745P
E
Bulb
dD214N
dK357N
dR364W
dW740S
µm
0
4
5µ
1µ
0.5µ
Protein domains
GD
Cat. site
HB1
Hinge
HB2

### Figure S6

**Figure S7. Mitochondrial phenotypes associated with CMT2A mutations in a *Marf*$^{KO}$ background.**
Confocal Airyscan image Z-projections (20 slices) of mitochondrial networks (mit::dendra2) from living motor neurons (dorsal clusters of ventral nerve cord) of male *Marf*$^{KO}$*/Y, 3xP3-mCherry* ; *OK371-GAL4, UAS-mit::dendra2* third instar larvae. Colour code: Z-depth in dorso-ventral axis. Intermediate panels: high magnification optic slices of mitochondria structures pinpointed by arrows on upper panels. Bottom panels: TEM mitochondria high magnification views in motor neurons (dorsal clusters) from larval ventral nerve cords. **(A)** *Marf*$^{KO}$ (no endogenous *Marf* expression). *Marf*$^{CTRL}$ (*UAS-HA::Marf*$^{WT}$). **(B)** Grape alleles (*UAS-HA::Marf*$^{CMT2A}$). **(C)** Tangle alleles (*UAS-HA::Marf*$^{CMT2A}$). **(D)** Bulb alleles (*UAS-HA::Marf*$^{CMT2A}$). **(E)** Peanut alleles (*UAS-HA::Marf*$^{CMT2A}$).

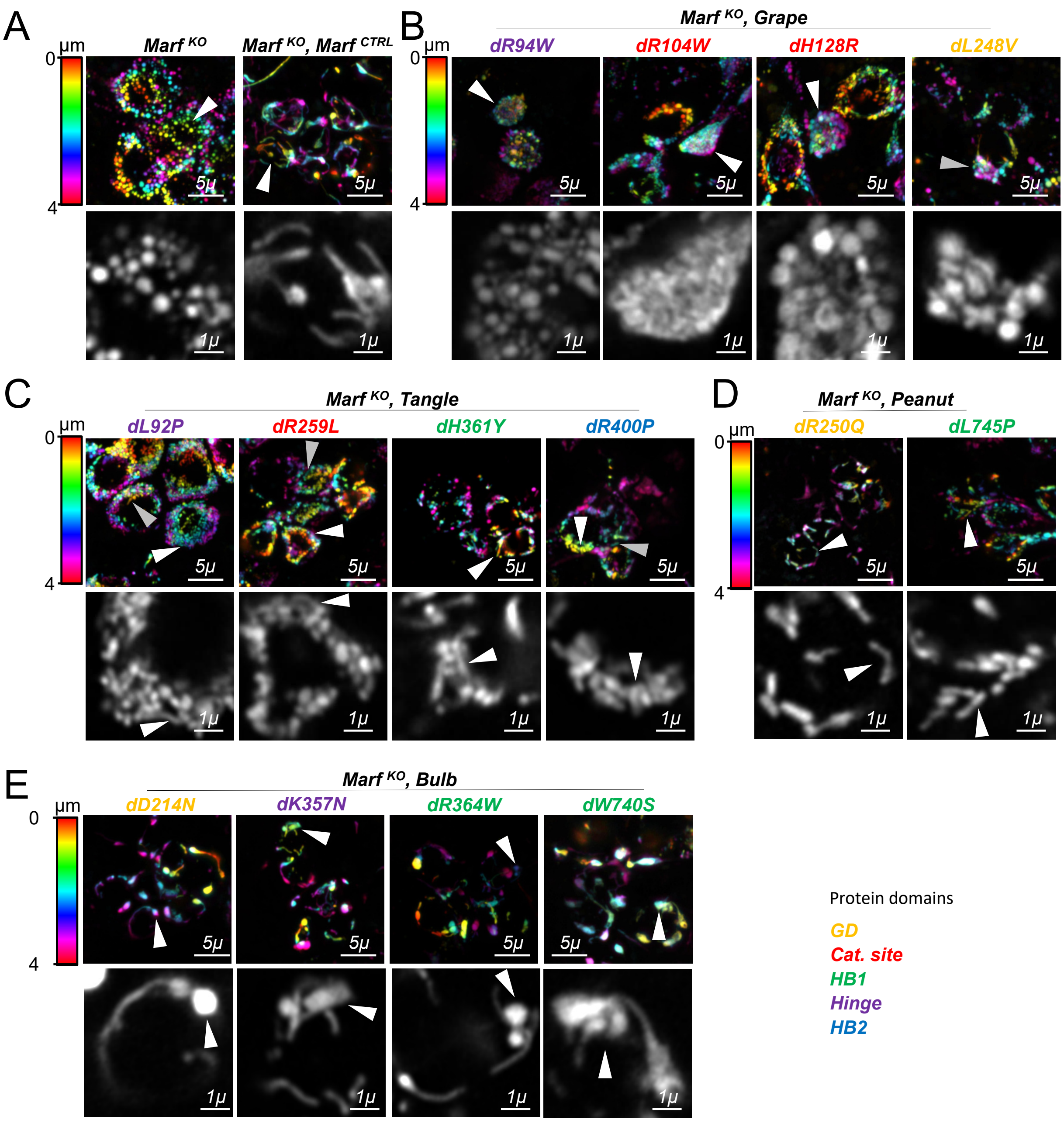
A
μm
0
4
Marf KO
Marf KO, Marf CTRL
5μ
1μ
B
Marf KO, Grape
dR94W
dR104W
dH128R
dL248V
C
Marf KO, Tangle
dL92P
dR259L
dH361Y
dR400P
D
Marf KO, Peanut
dR250Q
dL745P
E
Marf KO, Bulb
dD214N
dK357N
dR364W
dW740S
Protein domains
GD
Cat. site
HB1
Hinge
HB2

### Figure S7

**Figure S8. Mitochondrial morphology in Peanut mutants.**
Quantification of mitochondrial morphology in mit::dendra2-labelled motor neurons from larvae expressing only endogenous *Marf* (wild type), overexpressing wild-type *Marf* (*Marf* $^{CTRL}$) or Peanut *Marf* mutants *dR250Q* and *dL745P*. **(A)** Mean mitochondrial length +/- SD. MW U test: *versus* wild type p-value <0.05/*, p-value <0.001/***, *versus Marf* $^{CTRL}$ p-value <0.001/$^{\#\#\#}$. **(B)** Mean mitochondrial connectivity +/- SD (number of branches). MW U test: *versus* wild type p-value <0.001/***, *versus Marf* $^{CTRL}$ p-value <0.001/$^{\#\#\#}$. **(C)** Distribution of mitochondria according to their length. **(D)** Distribution of mitochondria according to their number of branches. For colour code see Figure 1F.

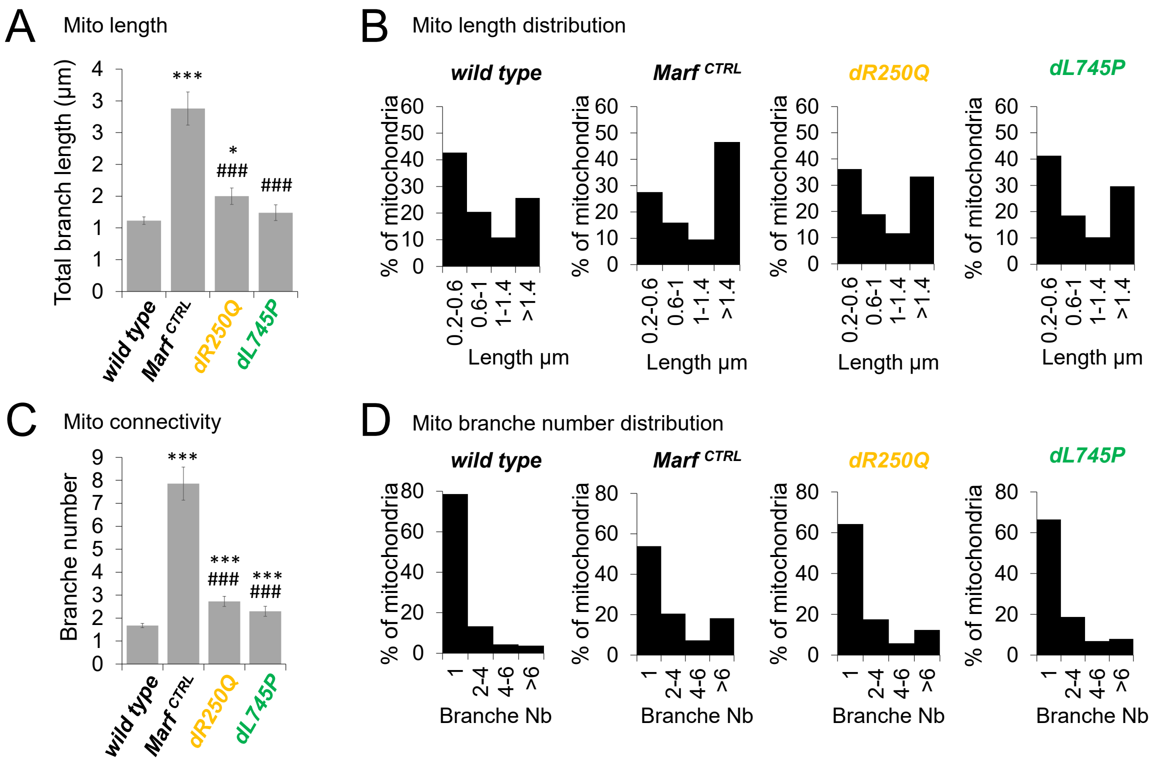
A
Mito length
Total branch length (µm)
wild type
Marf CTRL
dR250Q
dL745P
B
Mito length distribution
% of mitochondria
0.2-0.6
0.6-1
1-1.4
>1.4
Length µm
C
Mito connectivity
Branche number
D
Mito branche number distribution
1
2-4
4-6
>6
Branche Nb

**Figure S8**

**Figure S9. Startle-induced locomotion does not require vision.**

Average locomotor performance (startle-induced negative geotaxis) of 10 days old *elav-GAL4 ; UAS-mit::dendra2/+* adult flies placed in the dark or in the light. Histogram shows the proportion of flies according to the height they have reached (1: 0-15 mm, 2: >15-30 mm, dark 3: >30-45mm, 4: >45mm). Mean of three independent races each involving 10 flies.

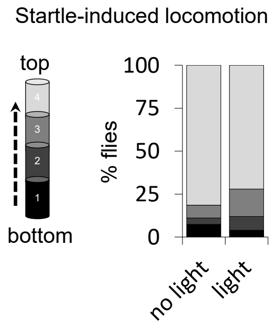
Startle-induced locomotion
top
4
3
2
1
bottom
100
75
50
25
0
% flies
no light
light

**Figure S9**

**Figure S10. Impact of CMT2A alleles on mitochondrial distribution, relative ER-mitochondrial localisation and mitochondrial NMJ content.**

**(A)** Confocal microscopy images of living motor neurons in *OK371-GAL4, UAS-mit::dendra2 ; UAS-myr::mRFP* third instar larva showing mitochondria and plasma membrane expressing *Marf* wild-type (*Marf* $^{CTRL}$) or mutant (*Marf* $^{CMT2A}$) transgenes from the four allelic groups. **(B)** Confocal Airyscan microscopy images of living motor neurons showing mitochondria and ER in *OK371-GAL4, UAS-mit::mKate2, UAS-KDEL::GFP* third instar larva expressing *Marf* wild-type (*Marf* $^{CTRL}$) or mutant (*Marf* $^{CMT2A}$) transgenes from the four allelic groups. Magnification views: ER and mitochondria in close vicinity and overlapping (arrow heads). **(C)** Confocal Airyscan microscopy images showing mitochondria within NMJ boutons at the surface of living muscles from *OK371-GAL4, UAS-mit::dendra2/UAS-HA::Marf ; UAS-myr::mRFP* third instar larva expressing *Marf* wild-type (*Marf* $^{CTRL}$) or mutant (*Marf* $^{CMT2A}$) transgenes from the four allelic groups.

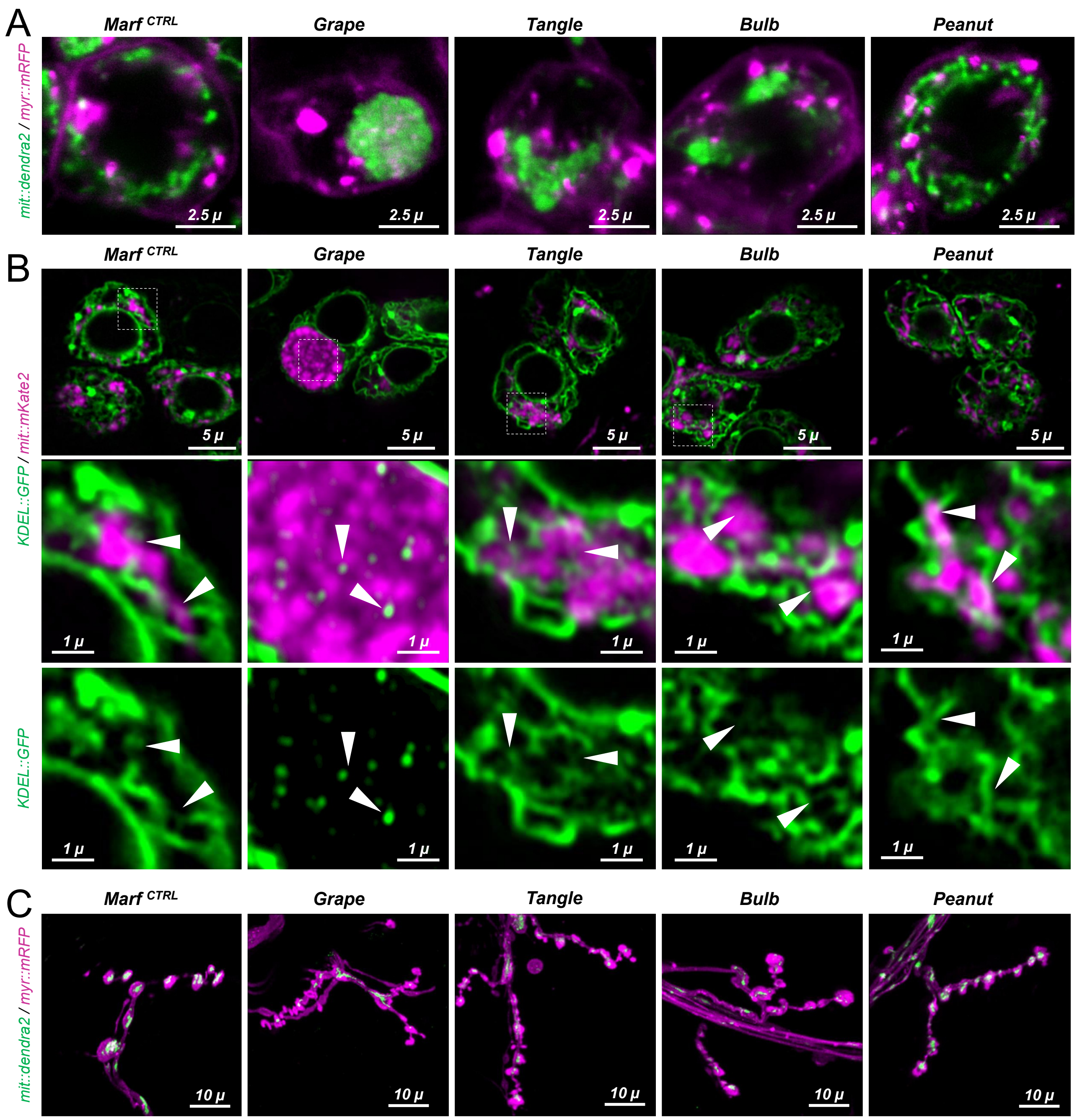
A
Marf CTRL
Grape
Tangle
Bulb
Peanut
mit::dendra2 / myr::mRFP
2.5 µ
B
5 µ
KDEL::GFP / mit::mKate2
1 µ
KDEL::GFP
C
10 µ
mit::dendra2 / myr::mRFP

**Figure S10**

**Figure S11. Parkin overexpression in CMT2A mutants.**

Confocal Airyscan image Z-projections (20 slices) of mitochondrial networks (mit::dendra2) from living motor neurons (dorsal clusters of ventral nerve cord) of wild type and CMT2A mutant *OK371-GAL4, UAS-mit::dendra2* third instar larvae. Colour code: Z-depth in dorso-ventral axis. Intermediate panels: high magnification optic slices of mitochondria structures pinpointed by arrows on upper panels. Bottom panels: TEM mitochondria high magnification views in motor neurons (dorsal clusters) from larval ventral nerve cords. **(A)** *Marf* $^{CTRL}$ (*UAS-HA::Marf* $^{WT}$). *Marf* $^{CTRL}$, *parkin*$^{OE}$ (*UAS-HA::Marf* $^{WT}$, *UAS-parkin*). *Marf* $^{CTRL}$, *parkin*$^{KD}$ (*UAS-HA::Marf* $^{WT}$ ; *UAS-parkin RNAi*). **(B)** Grape alleles, *parkin*$^{OE}$ (*UAS-HA::Marf*$^{CMT2A}$, *UAS-parkin*). **(C)** Tangle alleles, *parkin*$^{OE}$ (*UAS-HA::Marf* $^{CMT2A}$, *UAS-parkin*). **(D)** Bulb alleles, *parkin*$^{OE}$ (*UAS-HA::Marf* $^{CMT2A}$, *UAS-parkin*). **(E)** Peanut alleles, *parkin*$^{OE}$ (*UAS-HA::Marf*$^{CMT2A}$, *UAS-parkin*).

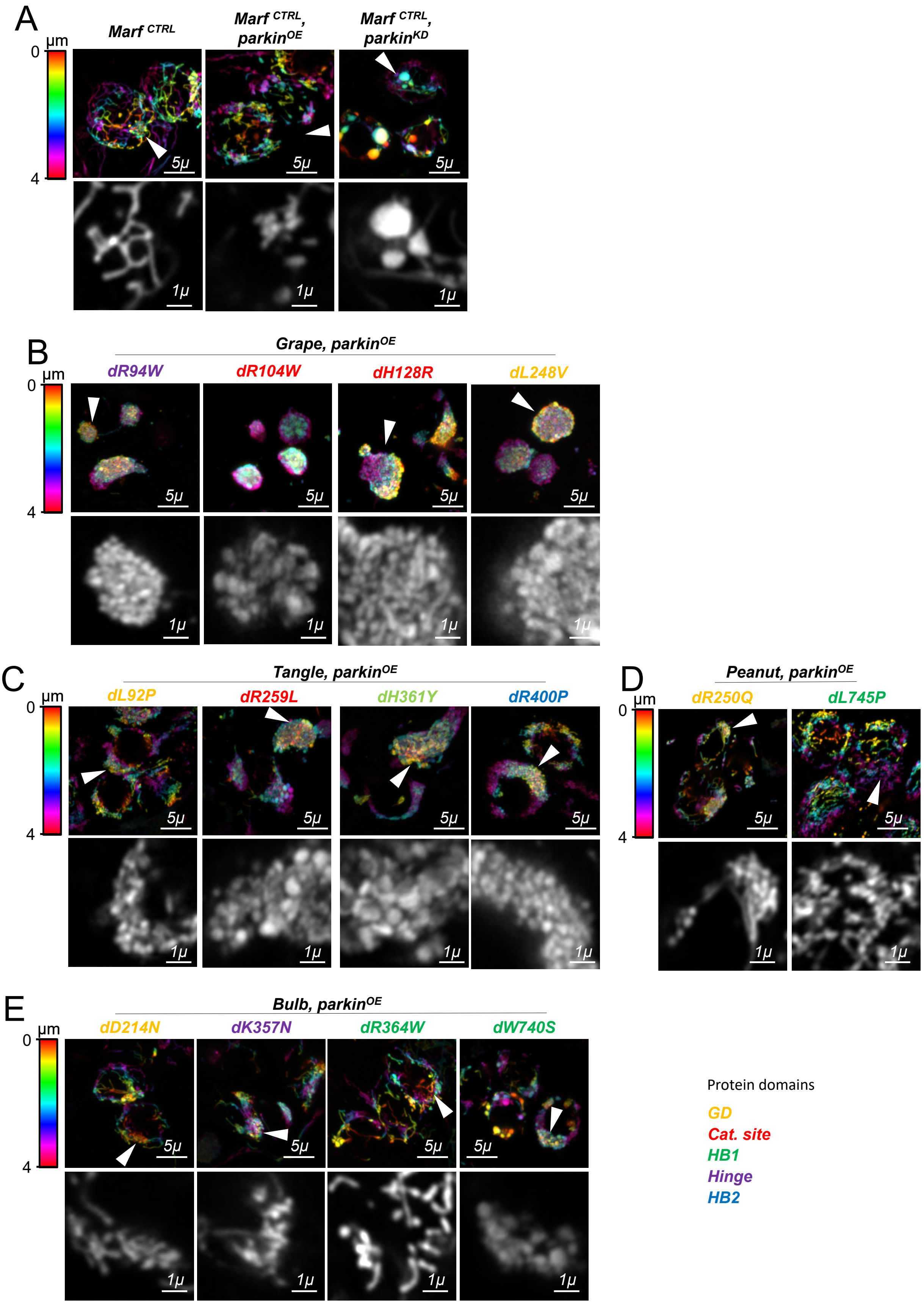
A
Marf CTRL
Marf CTRL, parkinOE
Marf CTRL, parkinKD
µm
0
4
5µ
1µ
B
Grape, parkinOE
dR94W
dR104W
dH128R
dL248V
C
Tangle, parkinOE
dL92P
dR259L
dH361Y
dR400P
D
Peanut, parkinOE
dR250Q
dL745P
E
Bulb, parkinOE
dD214N
dK357N
dR364W
dW740S
Protein domains
GD
Cat. site
HB1
Hinge
HB2

**Figure S11**

**Table S1. Correlation plots.**

**(A)** Correlogram showing Kendall coefficient (τ) between the indicated variables. **(B)** p-value corresponding to the level of significance of the correlations presented in A.

## A

| | | | | | Mito in wild type | | | | | | Mito in *Marf* $^{KO}$ | | | |
|---|---|---|---|---|---|---|---|---|---|---|---|---|---|---|
| | Locomotion | Distribution Mito | ER-Mito Overlap | NMJ Mito Content | % Fragmented | % Clustered | % Tubular | % Net-like | % Enlarged | Mito Length | % Fragmented | % Tubular | Ubi/Native Marf | Total Native Marf |
| Locomotion | 1 | 0.82 | 0.8 | 0.65 | -0.4 | -0.64 | 0.7 | 0.4 | 0.15 | 0.14 | -0.27 | 0.12 | 0.08 | -0.56 |
| Distribution Mito | | 1 | 0.76 | 0.65 | -0.45 | -0.59 | 0.63 | 0.45 | 0.12 | 0.19 | -0.32 | 0.12 | 0.12 | -0.56 |
| ER-Mito Overlap | | | 1 | 0.58 | -0.45 | -0.67 | 0.6 | 0.4 | 0.17 | 0.12 | -0.16 | 0.1 | 0.01 | -0.49 |
| NMJ Mito Content | | | | 1 | -0.31 | -0.59 | 0.5 | 0.35 | 0.15 | 0.23 | -0.32 | 0.12 | 0.21 | -0.56 |
| % Fragmented | | | | | 1 | 0.15 | -0.5 | -0.65 | -0.52 | -0.23 | 0.26 | -0.18 | -0.04 | 0.26 |
| % Clustered | | | | | | 1 | -0.44 | -0.16 | -0.29 | -0.35 | 0.43 | -0.35 | -0.32 | 0.61 |
| % Tubular | | | | | | | 1 | 0.41 | 0.35 | 0.1 | -0.18 | 0.1 | -0.05 | -0.38 |
| % Net-like | | | | | | | | 1 | 0.28 | 0.16 | -0.3 | 0.11 | 0.21 | -0.18 |
| % Enlarged | | | | | | | | | 1 | 0.46 | -0.38 | 0.44 | 0.25 | -0.36 |
| Mito Length | | | | | | | | | | 1 | -0.74 | 0.85 | 0.63 | -0.27 |
| % Fragmented | | | | | | | | | | | 1 | -0.67 | -0.58 | 0.32 |
| % Tubular | | | | | | | | | | | | 1 | 0.52 | -0.16 |
| Ubi/Native Marf | | | | | | | | | | | | | 1 | -0.38 |
| Total Native Marf | | | | | | | | | | | | | | 1 |

Row groups: Mito in wild type (% Fragmented to Mito Length); Mito in *Marf* $^{KO}$ (% Fragmented, % Tubular)

**Kendall Coefficient (τ)**

Scale: 1, 0.8, 0.6, 0.4, 0.2, 0, -0.2, -0.4, -0.6, -0.8, -1

## B

| | | | | | Mito in wild type | | | | | | Mito in *Marf* $^{KO}$ | | | |
|---|---|---|---|---|---|---|---|---|---|---|---|---|---|---|
| | Locomotion | Distribution-Mito | ER Mito Overlap | NMJ Mito Content | % Fragmented | % Clustered | % Tubular | % Net-like | % Enlarged | Mito Length | % Fragmented | % Tubular | Ubi/Native Marf | Total Native Marf |
| Locomotion | 1 | 0 | 1e-05 | 0.00076 | 0.05744 | 0.00333 | 0.00099 | 0.05854 | 0.50189 | 0.51825 | 0.19251 | 0.59064 | 0.74719 | 0.00457 |
| Distribution Mito | | 1 | 4e-05 | 0.00076 | 0.03315 | 0.00712 | 0.00329 | 0.03393 | 0.58272 | 0.38799 | 0.12679 | 0.59064 | 0.59064 | 0.00457 |
| ER-Mito Overlap | | | 1 | 0.00302 | 0.03315 | 0.00223 | 0.00477 | 0.05854 | 0.42742 | 0.59064 | 0.45056 | 0.66717 | 1 | 0.01384 |
| NMJ Mito Content | | | | 1 | 0.15004 | 0.00712 | 0.01869 | 0.09644 | 0.50189 | 0.27918 | 0.12679 | 0.59064 | 0.33081 | 0.00457 |
| % Fragmented | | | | | 1 | 0.52139 | 0.02638 | 0.00388 | 0.02506 | 0.27399 | 0.22663 | 0.38779 | 0.86287 | 0.22663 |
| % Clustered | | | | | | 1 | 0.0576 | 0.48234 | 0.22138 | 0.11179 | 0.05033 | 0.11179 | 0.14214 | 0.0049 |
| % Tubular | | | | | | | 1 | 0.06533 | 0.13274 | 0.63813 | 0.41048 | 0.63813 | 0.81409 | 0.07778 |
| % Net-like | | | | | | | | 1 | 0.22572 | 0.45616 | 0.15184 | 0.60592 | 0.32982 | 0.38988 |
| % Enlarged | | | | | | | | | 1 | 0.03263 | 0.07667 | 0.04395 | 0.24609 | 0.0993 |
| Mito Length | | | | | | | | | | 1 | 8e-05 | 0 | 0.00123 | 0.19251 |
| % Fragmented | | | | | | | | | | | 1 | 0.00045 | 0.00302 | 0.12679 |
| % Tubular | | | | | | | | | | | | 1 | 0.00975 | 0.45056 |
| Ubi/Native Marf | | | | | | | | | | | | | 1 | 0.06166 |
| Total Native Marf | | | | | | | | | | | | | | 1 |

Row groups: Mito in wild type (% Fragmented to Mito Length); Mito in *Marf* $^{KO}$ (% Fragmented, % Tubular)

**p-value**

- <0,001
- <0.001<0.01
- <0.01<0.05
- >0.05

### Table S1
